# Detecting and quantifying clonal selection in somatic stem cells

**DOI:** 10.1101/2021.12.15.472780

**Authors:** Verena Körber, Naser Ansari-Pour, Niels Asger Jakobsen, Rachel Moore, Nina Claudino, Marlen Metzner, Franziska Hörsch, Batchimeg Usukhbayar, Mirian Angulo Salazar, Simon Newman, Benjamin JL Kendrick, Adrian H Taylor, Rasheed Afinowi-Luitz, Roger Gundle, Bridget Watkins, Kim Wheway, Debra Beazley, Stephanie G Dakin, Andrew J Carr, Paresh Vyas, Thomas Höfer

## Abstract

As DNA variants accumulate in somatic stem cells, become selected or evolve neutrally, they may ultimately alter tissue function. When, and how, selection occurs in homeostatic tissues is incompletely understood. Here, we introduce SCIFER, a scalable method that identifies selection in an individual tissue, without requiring knowledge of the underlying driver event. Moreover, SCIFER infers the self-renewal and mutation dynamics of the tissue’s stem cells, and, if selection is present, the size and growth rate of the largest selected clone. We benchmark SCIFER with published data and then probe bone marrow of 22 non-leukemic individuals for clonal hematopoiesis (CH), identifying CH with known and unknown driver events. Unexpectedly, we find accelerated division of all stem cells in CH, compared to age-matched non-CH individuals, suggesting that the bone marrow environment alters stem cell dynamics in individuals with CH. SCIFER is broadly applicable to renewing somatic tissues to detect and quantify selection.

## Main

In phenotypically normal stem cells, somatic variation accumulates throughout development ^1^ and post-natal life ^2-6^ through diverse genetic mechanisms^7,8^. Somatic variants may alter stem cell function, with the potential to ultimately change tissue function. Over time, stem cell clones vary in size due to genetic drift^9,10^, or selection^4-6^. Understanding how somatic mosaicism is shaped by selection and genetic drift, and quantifying the underlying proliferation, differentiation and mutation dynamics of tissue stem cells, are fundamental questions.

Hematopoiesis is a paradigm for somatic mosaicism. In elderly individuals, clonal expansions occur in the hematopoietic stem cell (HSC) population^11-16^. This clonal hematopoiesis (CH) is associated with a heightened risk of developing blood cancer^12,13,15,17^, excess cardiovascular mortality^18^ and chronic infection^19,20^. Interestingly, a recent study suggests that atherosclerosis can drive CH by accelerating HSC proliferation and hence mutation acquisition^21^. Typically, CH is detected by probing blood samples for recurrent mutations in genes associated with leukemia, such as *DNMT3A, TET2* and *ASXL1*. More recently, blood cell clones, with known driver mutations, have been characterized in longitudinal studies^22,23^. When in life CH clones are born, and under which conditions driver mutations are generated and selected, remain key open questions.

Mathematical analyses of driver mutation patterns in large cohorts have shown that clonal expansion in CH occurs through selection^24^ and predict that clonal selection of HSCs is far more prevalent in humans than suggested by the observed number of CH cases with known drivers^25^. Hence, a large fraction of CH cases and their drivers may remain unidentified.Indeed, recent studies of human hematopoiesis found frequently expanded clones in elderly individuals for which the driver event was unknown^22,26^. Here, clonal selection was detected by sequencing 3050 whole genomes of single hematopoietic stem and progenitor cells (HSPCs) expanded ex vivo, from 11 donors; phylogenetic trees were reconstructed using somatic variants in these cells^2^.

These seminal observations raise the following questions: (a) Do CH drivers emerge by chance during normal HSC turnover, or does increased turnover or/and somatic mutation rate facilitate acquisition of driver events that promote selection? (b) Do stem cell clones with known CH driver genes, and clones with unknown drivers, differ in their frequency in the human population and in their clinical phenotypes? (c) When do selection events, with known or unknown drivers, occur in life, and (d) how fast do the selected clones grow with known and unknown driver events? These questions are more easily addressed with a more cost effective, scalable method, than phylogenetic tree reconstruction, to identify clonal selection and quantitate stem cell dynamics. Ideally, such a method would also work in tissues where stem cells cannot be cultured ex vivo. A recent approach used WGS of single blood samples to search for CH, exploiting the fact that selection of a mutated HSC will often increase the total count of somatic variants detectable by WGS^27^. This method uses prior knowledge of driver mutations and cannot reconstruct the underlying evolutionary dynamics.

Here, we show that neutral evolution and clonal selection leave characteristic imprints in the statistics of somatic variants detected by bulk WGS at a single snapshot, irrespective of whether the driver of selection is known or not. Using this observation, we develop population-genetics theory for the accumulation and spread of somatic variants in development and subsequent homeostatic tissue renewal, either in the absence, or presence, of selection. Based on this theory, we devise a Bayesian inference framework that discriminates clonal expansion due to selection from neutral evolution, and quantifies the underlying stem cell parameters in a homeostatic tissue. We validate our method with synthetic data and published experimental data, and then apply it to a new cohort of bone marrow samples of non-leukemic individuals.

## Results

### Quantifying clonal selection in somatic mosaics

The vast majority of genetic variants acquired in somatic stem cells are neutral^28^. We reasoned that the quantitative distribution of these variants measured in stem cells of an individual provides information on the underlying dynamics of genetic drift and selection. To derive the expected distribution and subsequently compare it to experimental data, we considered developmental expansion of a renewing tissue, such as the blood, the skin and the gut epithelium, followed by a phase of homeostatic turnover. During development, stem cell divisions feed tissue expansion, whilst in adulthood stem cell self-renewal usually balances loss, by differentiation and cell death (Fig. 1a and Methods for a mathematical description of the model). We focussed specifically on the abundant somatic single-nucleotide variants (SSNVs) acquired during both development and homeostasis and inherited to progeny. As the number of developing tissue stem cells is initially small, few SSNVs acquired early in life (Fig 1b; variants A and B); but they will have high variant allele frequencies (VAFs). Later, when the stem cell numbers are larger, more variants are acquired in total but now each variant will be at a lower VAF (Fig 1b; variants C, D and E). Owing to the inherent stochasticity of self-renewing divisions and loss of stem cells, neutral SSNVs will drift with time so that some variants will randomly increase their VAF (Fig. 1b; variants B, C, D and F) while others either decrease, or are extinguished (Fig. 1b; variant E). To quantify the acquisition of SSNVs and the changes in their frequencies, we model the neutral clonal dynamics during development and homeostasis by combining expanding (supercritical) and homeostatic (critical) birth-death processes (see Methods, Population-genetics model). We find an analytical solution for the VAF distribution as a function of time that lends itself to parameter inference from experimental data.

**Fig. 1.**
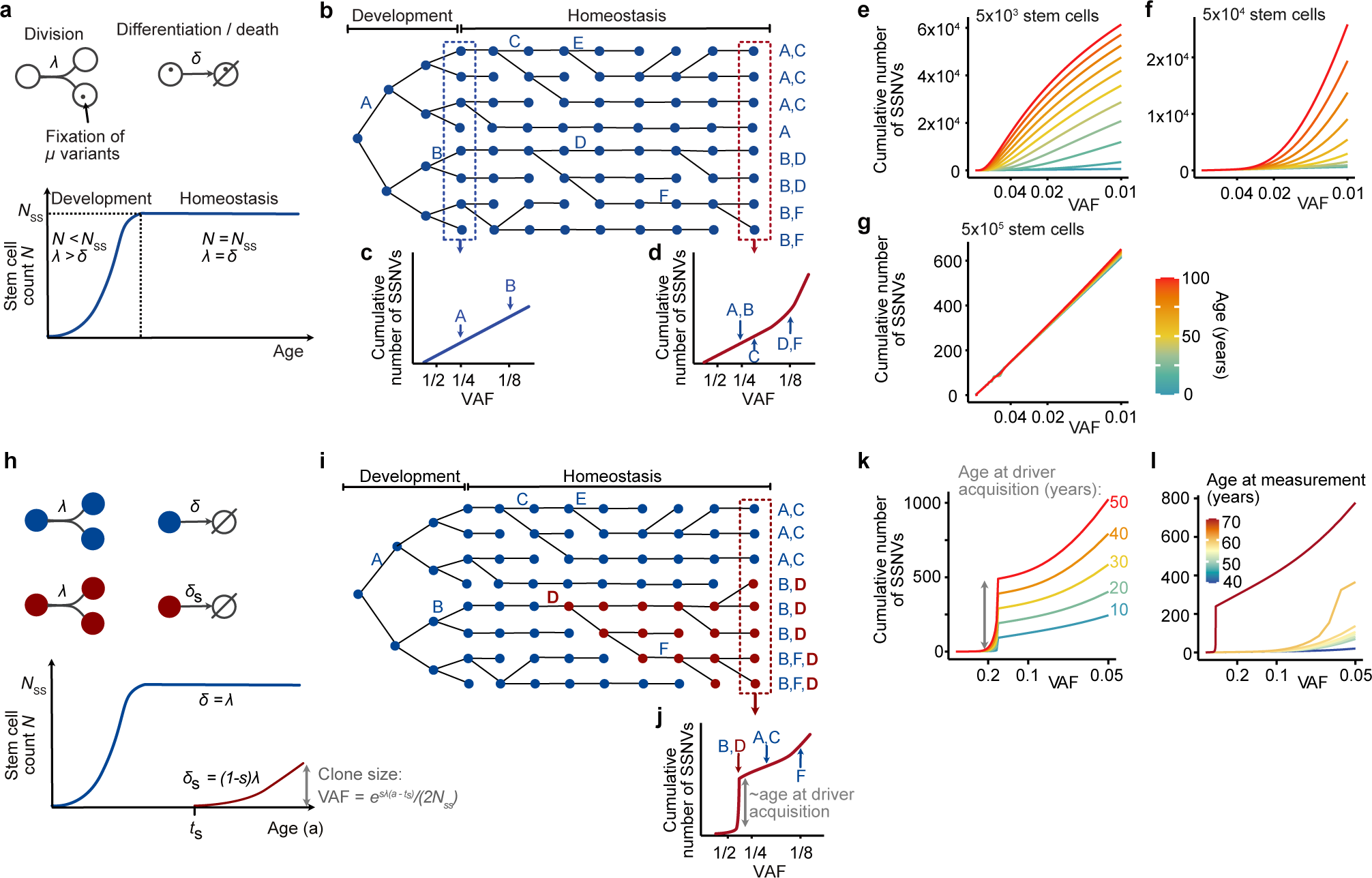
Population-genetics model of drift and selection in homeostatic tissues. **a**, Modelled processes and associated parameters in the model of drift. Stem cells either divide symmetrically with rate λ, or exit the stem cell compartment by differentiating (or dying), with rate *δ*. The blood stem cell count (*N*) increases in development (λ > *δ*) until reaching steady state numbers (*N*_*ss*_) and remains constant during adulthood (*N* =*N*_ss_ and λ = *δ*). On average, cells acquire μ neutral variants during each cell division. **b**, Schematic illustrating variant accumulation during development and subsequent homeostasis. **c**, Cumulative number of somatic single nucleotide variants (SSNV) versus variant allele frequency (VAF) in development (where the scaling of the x-axis is transformed to 1/VAF to spread out low-frequency variants). **d**, Cumulative number of SSNV versus VAF in adult life (where the scaling of the x-axis is transformed to 1/VAF to spread out low-frequency variants). **e**, Simulated cumulative VAF distribution of SSNVs at selected ages between 0 and 100 years for 5x10^3^ stem cells; λ=5/year, μ=10/division. **f**, Simulated cumulative VAF distribution of SSNVs at selected ages between 0 and 100 years for 5x10^4^ stem cells; λ=5/year, μ=10/division. **g**, Simulated cumulative VAF distribution of somatic SSNVs at selected ages between 0 and 100 years for 5x10^5^ stem cells; λ=5/year, μ=10/division. **h**, Model of clonal selection. A selective driver event reduces the loss rate (differentiation or death) by a factor *s*, causing selective outgrowth of the mutant clone (red); the remaining parameters are defined in **a**. The VAF of the selected clone increases exponentially with the age at measurement, *a*. **i**, As in **b**, but here an acquired driver mutation (D) causes the selective outgrowth of the mutant clone (red). **j**, All variants in the selected clone’s cell of origin are inherited to its progeny and hence reach a high VAF during clonal expansion, reflected in a shoulder in the cumulative VAF distribution. **k**, Simulated cumulative VAF distributions when a driver mutation is acquired at different ages, and the SSNVs are measured 45 years later, when the clone has reached a size of 32%. Note that, the later in life the selected clone is born, the more somatic variants have accumulated in its founder cell, and hence the larger the height of the shoulder. In the simulation, the selected clone grows by 22% per year (*s=*0.02, λ=10/year, μ=1/division, *N*_ss_=25,000). **l**, Simulated cumulative VAF distributions measured at varying ages after a driver mutation was acquired at 20 years of age. As the time interval between the acquisition of the selective driver mutation and the sampling of the tissue increases, the selected clone will be larger and eventually reach fixation. Therefore, SSNVs defining the founder cell of the selected clone will have higher VAFs, and the position of the shoulder on the x-axis will progressively move to the left. As in **k**, the selected clone grows by 22% per year (*s=*0.02, λ=10/year, μ=1/division, *N*_ss_=25,000).

How do the two principal phases – development and homeostasis – shape the VAF distribution measured in adulthood? During developmental expansion, genetic drift is negligible irrespective of the total number of stem cells, resulting in an invariant linear relation between cumulative SSNV count and 1/VAF (Fig. 1c; to zoom in on small VAFs, we plot the cumulative SSNV distribution against 1/VAF)^29,30^. During homeostasis, genetic drift might alter the cumulative VAF histogram by increasing the number of SSNVs with low VAF (Fig.1d). The extent of homeostatic drift depends on the number of stem cells and their self-renewal rate. If there are 5x10^3^ stem cells dividing 5 times per year, drift will increase the variant count with high VAF over the human lifespan (Fig. 1e; given that the development of HSCs and major increase in number occurs in the fetus, in the model we assumed transition from development to approximate homeostasis around birth). For 5x10^4^ stem cells and the same division rate, the variant count will increase with age at lower VAFs (right part of the VAF distribution) and remain stable at high VAFs (left part of the VAF distribution) (Fig. 1f). Finally, with 5x10^5^ stem cells, the distribution is nearly stable with age for all VAFs >1% (Fig. 1g). Prior work has inferred, for a 59-year old-human, that 4x10^4^ -2x10^5^ HSCs divide 0.6 -6 times per year^2^. With these values, our model suggests the VAF histogram at high VAF (about 10% or higher) is insensitive to genetic drift over the human lifespan but might be modified by drift at low VAF.

Next, we modelled how clonal selection alters the VAF distribution. We consider the leading selected clone, born at time *t*_s_ consequent to a driver event with selective advantage *s*, promotes mutant clone expansion and concomitant normal stem cell displacement (Fig. 1h and i). With clonal selection, neutral variants may have originated: (i) in normal stem cells before acquisition of the driver event; (ii) within the selected clone or (iii) in non-selected stem cells after the driver event occurred. We find analytical expressions for the evolution of all three types of variants, over time, which together yield the cumulative VAF distribution of SSNVs within a tissue, in the presence of a selected clone (Methods). Expansion of a selected clone will increase the VAF of the founding driver mutation (mutation D in Fig. 1i) and of all other neutral SSNVs in the founding cell of the selected clone. As a consequence, the selected clone generates a shoulder in the cumulative VAF distribution (Fig. 1j), as opposed to the gradual increase of the cumulative SSNV number in the case of neutral evolution (Fig. 1d). Importantly, the shape of the distribution discriminates between selection and neutral evolution, whereas the mere variant count detectable in WGS may fail to do so (compare the VAF histograms in Fig. 1j, shaped by selection, with Fig. 1e-f, shaped by drift). Hence, the shoulder in the cumulative VAF distribution provides a robust signature of clonal selection agnostic of the identity of the driver.

The height of the shoulder increases with the age at which the selected clone originated, *t*_s_, (Fig. 1k), since the number of neutral variants in the clone’s stem cell of origin is the higher, the older the individual as more prior cell divisions have occurred. As the selected clone expands with time, the shoulder reaches higher VAFs (i.e., the shoulder moves to the left on the x-axis the greater the time interval between the birth of the clone and measurement of SNNVs and their VAFS by WGS; Fig. 1l). The VAF reached in the time interval between clone origin and sample acquisition yields the selective advantage *s* (Fig. 1h, lower panel). Thus, height and position of the shoulder allow inference of the clone’s age and selective advantage.

### Selection detected in noisy WGS data

WGS measures the VAF of a variant with an error that is approximately binomially distributed, with the variance depending on sequencing depth^31^. To understand how this measurement error influences the ability to identify clonal selection in bulk WGS data, we simulated WGS data with our models of neutral evolution (Fig. 1a,b) and selection (Fig. 1h,i). Briefly, we performed repeated stochastic simulations of the two models, generating 10 data sets with neutral evolution and 70 data sets with selection (with a driver mutation acquired at 20 years of age causing stochastic clonal expansion with an average growth rate of 22% per year). To account for the WGS measurement error, we drew variant reads from binomial distributions for average sequencing depths of 30x, 90x or 270x (Methods; Supplementary Table 1). We then used approximate Bayesian computation (ABC) to fit both the model of neutral evolution and the selection model to each data set (Extended Data Fig. 1), yielding posterior probabilities for the neutral model and the selection model to reproduce the noisy data.

To determine sensitivity and specificity in detecting selection, we used the posteriors to compute receiver-operating curves (ROC) for identifying clones of different sizes. At 90x sequencing depth, we found that selected clones with VAF ≥ 5% were detected reliably (Fig. 2a; area under the curve, AUC = 0.88 for VAF = 5% and AUC ∼ 1 for VAF = 7.5%), whereas smaller clones were not (Fig. 2a; VAF = 2.5%; AUC = 0.5). Discrimination between selection and neutral evolution was optimal when using a threshold of 15% for the posterior probability of the selection model, such that a posterior >15% identified selection (Fig. 2a, the operating points correspond to the 15% threshold, achieving ∼100% sensitivity and 80% specificity for clone size 5% VAF and ∼100% sensitivity and specificity for clone size 7.5% VAF).Exemplary selection posteriors for simulated 90x WGS data sets show discrimination between neutral evolution (0% VAF) and selection of clones ≥5% VAF (Fig. 2b). Further, our ROC analysis predicts that 30x WGS will allow detection of large selected clones > 20% VAF, while deeper WGS will enable detection of smaller clones, with VAF > 1% at 270x (Fig. 2c). Taken together, our method for **s**elected-**c**lone **i**n**fer**ence, termed SCIFER, is suitable for detecting clonal selection in bulk WGS data.

**Fig. 2.**
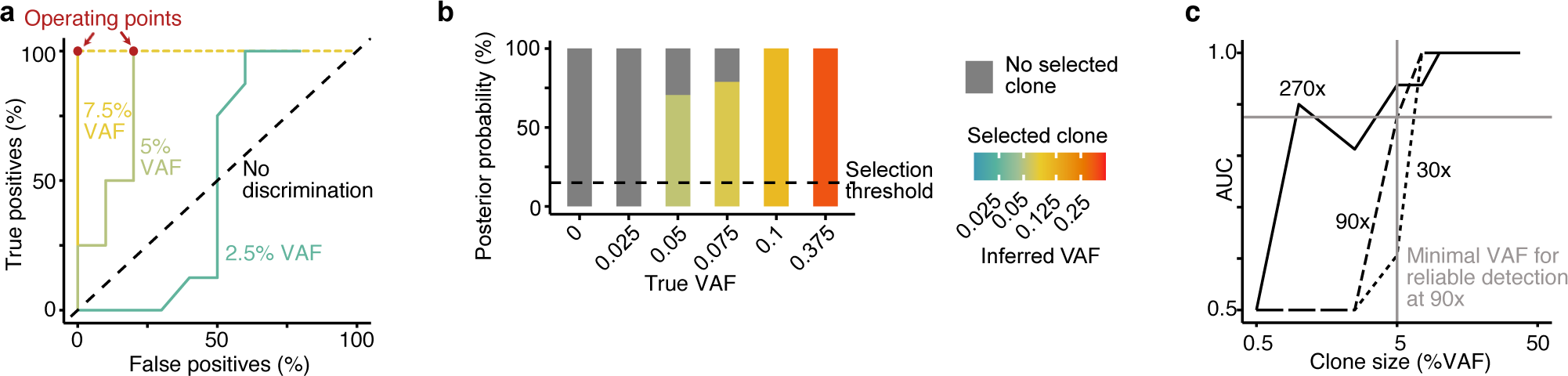
Benchmarking SCIFER with simulated data. **a**, Receiver-operating curves (ROCs) quantifying the ability of SCIFER to accurately detect clonal selection for different sizes of a selected clone (color encoded). ROCs were generated by applying SCIFER to simulated data, generated with a stochastic birth-death process with (s=0.02, corresponding to a selective advantage of 2% increase in birth versus death) or without (s=0) selection of a clone initiated at 20 years-of life (λ=10/year, μ=1/division, *N*_ss_=25,000, and assuming sequencing with an average coverage of 90x). In total 63 cases with selected clone sizes of VAF 0%, 2.5%, 5%, 7.5%, 10%, 25% and 37.5%, were generated. Models with and without clonal selection were fit to the data using approximate Bayesian computation. True positives and false positives were evaluated for varying posterior probability thresholds of clonal selection. For selected clones with VAFs ≥5%, the difference between true positives and false positives was maximal for a selection threshold of 15% (operating points, shown in red). **b**, Posterior probability for clonal selection (colored bars) and neutral evolution (grey bars) conditioned on selected clones with VAF ≥5% for six simulated cases (STN8, STS25, STS39, STS46, STS53, STS84) with varying clone size. Shown are true and inferred VAFs of the selected clone. The dashed line marks the selection threshold at 15% conditional posterior probability. **c**, Accuracy of SCIFER to distinguish clonal selection from genetic drift in simulated whole genome sequencing data. Shown are areas under the curve (AUC) computed from the receiver-operating curves (ROCs) shown in (**a**) and from ROCs obtained in analogy for simulated sequencing depths of 30x and 270x. The simulated data were generated as in (**a**). For sequencing depths of 30x and 90x, 63 cases with selected clone sizes of VAF 0%, 2.5%, 5%, 7.5%, 10%, 25% and 37.5%, were generated. For 270x sequencing depth, an additional 17 cases with selected clone sizes of 0.5% VAF and 1% VAF were used for model evaluation.

### Benchmarking SCIFER with phylogenetic trees of human hematopoiesis

Next, we benchmarked SCIFER with published WGS data from human HSPC single-cell clones, where the absence or presence of expanded clones is visible in reconstructed phylogenetic trees^2,22,26^. For a 59-year-old human, where hematopoiesis evolved neutrally, key HSC parameters (HSC number, self-renewal rate and the rate of acquiring somatic variants) were inferred based on WGS of 140 HSPC clones and targeted sequencing of peripheral blood cell populations^2^. This provided a quantitative reference for SCIFER. In addition to the experimental error in VAF measurement (cf. Extended Data Fig. 1), a further source of error in WGS is the calling of genome-wide somatic variants. Therefore, we used a stringent calling approach for identifying low-VAF variants in bulk WGS, intersecting the results of Mutect2 and Strelka^32^ (Extended data Fig. 2a). To examine the robustness of SCIFER with respect to variant calling, we also performed our analyses on the original variant calls obtained with Caveman^2^, which includes a substantial number of additional SSNVs (Extended data Fig. 2b-c). First, the reconstruction of the phylogenetic tree using our calling approach recapitulated the original work, yielding long branches with coalescent events located mainly at the top of the tree (Fig. 3a). Second, we generated pseudo-bulk WGS data from the single-cell variants and used SCIFER to fit the cumulative VAF distribution (Fig. 3b; Extended Data Fig. 2d). Irrespective of the calling algorithm, and in agreement with the original work, SCIFER inferred neutral evolution (Fig. 3c). Third, SCIFER inferred HSC number (Fig. 3d) and rate of self-renewing divisions (Fig. 3e), with close estimates for the two variant callers (6x10^4^-2x10^5^ HSCs dividing 3 to 8 times per year using the variants from Caveman; 2x10^4^-5x10^4^ HSCs dividing 0.4 to 6 times per year using the variants from Mutect2/Strelka; ranges are 80% credible intervals), which both overlapped with the estimate of the original work (4x10^4^-2x10^5^ HSCs dividing 0.6 to 6 times per year) ^2^. Finally, the rate of SSNV acquisition inferred by SCIFER (3-5 SSNVs/cell division) (Fig. 3f) overlapped with the estimate in the original work (3-28 SSNVs per division) when using the variants from Caveman. As expected, this rate was smaller (0.6-1 SSNV per division) with our conservative variant calls using the intersect of Mutect2 and Strelka. Interestingly, SCIFER could infer HSC number, self-renewal rate and mutation rate separately (Extended Data Fig. 2e, f). This feature relies on our use of absolute variant counts for each individual, as many rapidly dividing stem cells accumulate more variants in total than few rarely dividing stem cells (although the ratio of stem cell number and self-renewal rate may remain unchanged^2,24^), and the presence of both expansion and homeostatic phases (Supplementary Note 1 and Supplementary Fig. 1). In conclusion, SCIFER detected neutral evolution and quantified HSC number and division rate from a single pseudo-bulk sample. The inference was robust with respect to the variant calling algorithm, indicating that key aspects of stem cell dynamics can be robustly inferred from the VAF histogram.

**Fig. 3.**
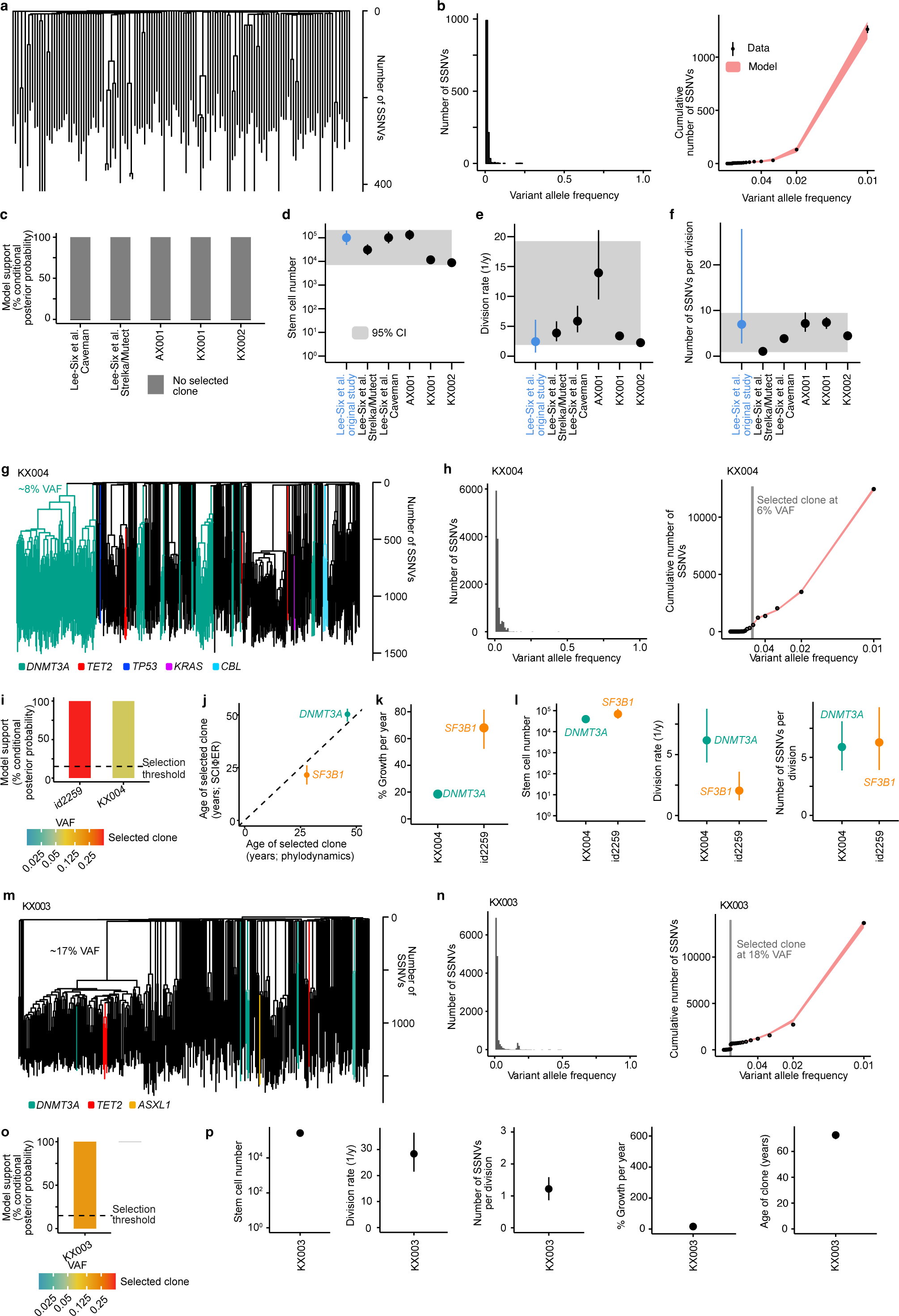
Benchmarking SCIFER with published pseudobulk data. **a**, Reconstructed single-cell phylogenies after re-calling SSNVs and indels from single-cell WGS data^2^. **b**, Left, VAF distribution of SSNVs shown in **a**, truncated at 1%. Right, model fit to the cumulative 1/VAF distribution (points and error bars, mean and standard deviation of the measured data; red area, 95% posterior probabilities of the model fit). **c**, Posterior probability for neutral evolution for pseudo-bulk WGS data from^2^ (labelled Lee-Six) and three samples (AX001, KX001 and KX002) from^26^; SCIFER did not support selection for a clone size >5% VAF. SCIFER was applied twice to the data from Lee-Six et al.^2^, using the SSNV counts obtained with Caveman or with the intersection of Mutect and Strelka. **d**, Inferred hematopoietic stem cell number (left), division rate (middle) and mutation rate (right) for the cases shown in **c** (median and 80% credible intervals for each sample; grey areas, 95% confidence band for all samples). Estimates of these parameters from^2^ are given for comparison. **e**, Single-cell phylogeny of published sample KX004^26^. Clones with variants in *DNMT3A* (green), *TET2* (red), *KRAS* (purple) and *CBL* (cyan) are shown. **f**, Left, VAF distribution of SSNVs of sample shown in **e**, truncated at 1% VAF. Right, model fit to the cumulative 1/VAF distribution (points and error bars, mean and standard deviation of the measured data; red area, 95% posterior probabilities of the model fit; grey area, 80% credible interval of the estimated clone size). **g**, Posterior probabilities for selection (conditioned on clones with VAF ≥5%) and neutral evolution for pseudo-bulk WGS data of two samples: id2259 described in Extended Data Fig. 2g, with known CH driver mutations in *SF3B1* (id2259, ref.^*22*^) *CBL* and (id KX004, ref.^26^). The color scale shows the inferred VAF of selected clone. The dotted line shows the 15% selection threshold. **h**, Comparison of age of the selected clones in the cases shown in **g**, estimated by SCIFER and by phylodynamic modelling in the original publications (points, median; error bars, 80% credible intervals). **i**, Estimated growth rates of the selected clones in the samples shown in **g** (points, median; error bars, 80% credible intervals). **j**, Estimated hematopoietic stem cell number for the two cases shown in **g**. Shown are median and 80% credible interval. **k**, Estimated HSC division rate for the two cases shown in **g**. Shown are median and 80% credible interval. **l**, Estimated number of new SSNVs acquired per cell division for the two cases shown in **g**. Shown are median and 80% credible interval. **m**, Single-cell phylogenies in published sample KX003 (ref.^26^), where the large expanded clone had an unknown driver event. Small clones with variants in *DNMT3A* (green), *TET2* (red) and *ASXL1* (orange) are also shown. **n**, Left, VAF distribution of SSNVs shown in **m**, truncated at 1%. Right, model fit to the cumulative 1/VAF distribution (points and error bars, mean and standard deviation of the measured data; red area, 95% posterior probabilities of the model fit; grey area, 80% credible interval of the estimated clone size). **o**, Posterior probabilities for clonal selection (conditioned on clones with VAF ≥5%) for pseudo-bulk WGS data of sample KX003 (ref.^26^). The color scale shows the inferred VAF of selected clone. The dotted lines show the 15% selection threshold. **p**, Estimated stem cell parameters (hematopoietic stem cell number, stem cell division rate and number of new SSNVs acquired per cell division) and selection parameters (growth rate and age of selected clone) for the sample shown in **m**. Shown are median and 80% credible intervals.

To further validate SCIFER’s ability to identify neutral evolution, we generated pseudo-bulk data from three further neutrally evolving, published human HSPC single-cell WGS datasets (AX001, KX001 and KX002) for which separate estimates of HSC number and division rate were not available^26^ (Extended Data Fig. 2g,h). In all cases, SCIFER correctly detected neutral evolution (Fig. 3c) and inferred HSC parameters broadly consistent with the estimate for the data set from Lee-Six et al ^2^ (Fig. 3d,f). Interestingly, for the oldest individual (AX001, 63 years of age, compared to KX001 and KX002, 29 and 38 years of age, respectively), the inferred HSC division rate was higher than for the other individuals (Fig. 3e). These data further support SCIFER’s ability to detect and quantify neutral evolution.

Next, we asked whether SCIFER detects clonal selection. WGS data from 448 single HSPCs from two published samples KX0004^26^ and id2259^22^ was used to construct phylogenetic trees. In both samples, CH driver mutations had been identified. KX004 harbored a major clone with a *DNMT3A* mutation (∼8% VAF) and several smaller clones (Fig. 3g); id2259 had an expanded clone with a mutation in the splicing factor *SF3B1* (with a size of nearly 50% VAF) containing a small *CBL* subclone (Extended Data Fig. 2g). In pseudo-bulk WGS data generated from single-cell SSNVs for both samples, SCIFER detected the respective leading *DNMT3A* (VAF 6%) and *SF3B1* (VAF 49%) clones (Fig. 3h; Extended Data Fig. 2i). SCIFER also inferred that these clones had expanded through selection (Fig. 3i). Using coalescent theory, the original studies estimated that the *SF3B1* mutation arose 25-30 years, and the *DNMT3A* mutation 45-50 years, prior to tissue sampling. SCIFER determined very similar clone ages (Fig. 3j). The estimated clonal growth rates were 68% per year for the *SF3B1* mutation and 18% for the *DNMT3A* mutation (Fig. 3k), in line with previous results suggesting stronger selection of *SF3B1* mutant clones^23^. The inferred stem cell number, division rate and rate of acquired SSNV/cell division (Fig. 3l) were in the same range as estimates for individuals without CH (Fig. 3d-f). Finally, we applied SCIFER to a previously published sample, KX003^26^, where phylogenetic reconstruction showed a large expanded clone (∼17% VAF) that harbored no known CH driver and that contained smaller clones with known drivers (Fig. 3m). Applying SCIFER to WGS pseudo-bulk data indeed detected a leading clone of 18% VAF (Fig. 3n) and inferred selection (Fig. 3o). The quantitative parameters – HSC number, division rate, rate of acquired SSNV/cell division, clonal growth rate and age of the clone – were in the same ranges as in the two samples above with known CH driver mutations (Fig. 3p, compare with Fig. 3j-l).

In summary, SCIFER robustly identified and quantified neutral evolution and selection in pseudo-bulk WGS data of human HSPCs, without using prior knowledge of the presence of a driver.

### SCIFER uncovers complexity of somatic mosaicism in bulk samples

Having established the performance of SCIFER, we applied it to genuine bulk WGS data from 22 humans, 30-89 years of age (Supplementary Table 2). We used human bone marrow (BM) samples collected from individuals at total hip replacement surgery. The samples were screened for CH by targeted sequencing of DNA from bone marrow mononuclear cells using a gene panel covering 97 known driver genes sequenced at an average of 800x to detected mutations with a VAF of ≥1% VAF(Supplementary Table 3)^33^. All individuals had normal blood counts and blood films, without an antecedent history of blood or inflammatory or autoimmune disorder^33^.

We selected 10 individuals without known drivers and 12 individuals with at least one mutation in either *DNMT3A* (6 individuals), *TET2* (4 individuals) or *ASXL1* (2 individuals). 90-120x WGS was performed on FACS-sorted Lin^-^CD34^+^ HSPCs (Extended Data Fig. 3a-d). A comparison with WGS results for mature BM mononuclear cells is given in Supplementary Note 2 and Supplementary Fig. 2. 30x WGS from hair follicle DNA was used as a germline control. We called SSNVs, small indels, copy number variants (CNVs), and structural variants (SVs) (see Methods). The WGS data confirmed either the presence, or absence, of known CH driver mutations in each sample that was originally noted in panel sequencing.

Applying SCIFER to the WGS data of the 10 cases without known CH drivers, seven cases were classified (labelled N1-N7) as neutrally evolving. However, in the remaining 3 cases (U1-U3) SCIFER detected clonal selection, with unknown driver events. For the 12 individuals with known CH drivers, SCIFER detected a leading clone in 10 of them, that could be associated with the CH driver mutation found by panel sequencing (A1-A2, with *ASXL1* mutation; D1-D5, with *DNMT3A* mutation; T1-T3, with *TET2* mutation). In the remaining two samples (U4 and U5), SCIFER detected a larger additional clone, without a known CH driver, whilst the driver identified by panel sequencing was associated with a smaller and different clone. Hence, SCIFER uncovered complexity of somatic mosaicism beyond the insights offered by panel sequencing of CH drivers.

### Quantifying neutral evolution

We now report the SCIFER results in detail. We focus first on the seven cases (N1-N7) without known CH driver mutations where neutral evolution was inferred by SCIFER (Fig. 4a-b; Extended Data Fig. 5; 2 males and 5 females, ranging from 30 to 76 years of age). The somatic single-base-pair substitution profiles agreed with neutral mutation accumulation during normal ageing (Extended Data Fig. 4a)^2,26,34^. Moreover, we found neither copy number alterations (CNAs), nor structural variants (SVs) in driver genes in these samples (Extended Data Fig. 4b, Supplementary Tables 4 and 5), consistent with neutral evolution.

**Fig. 4.**
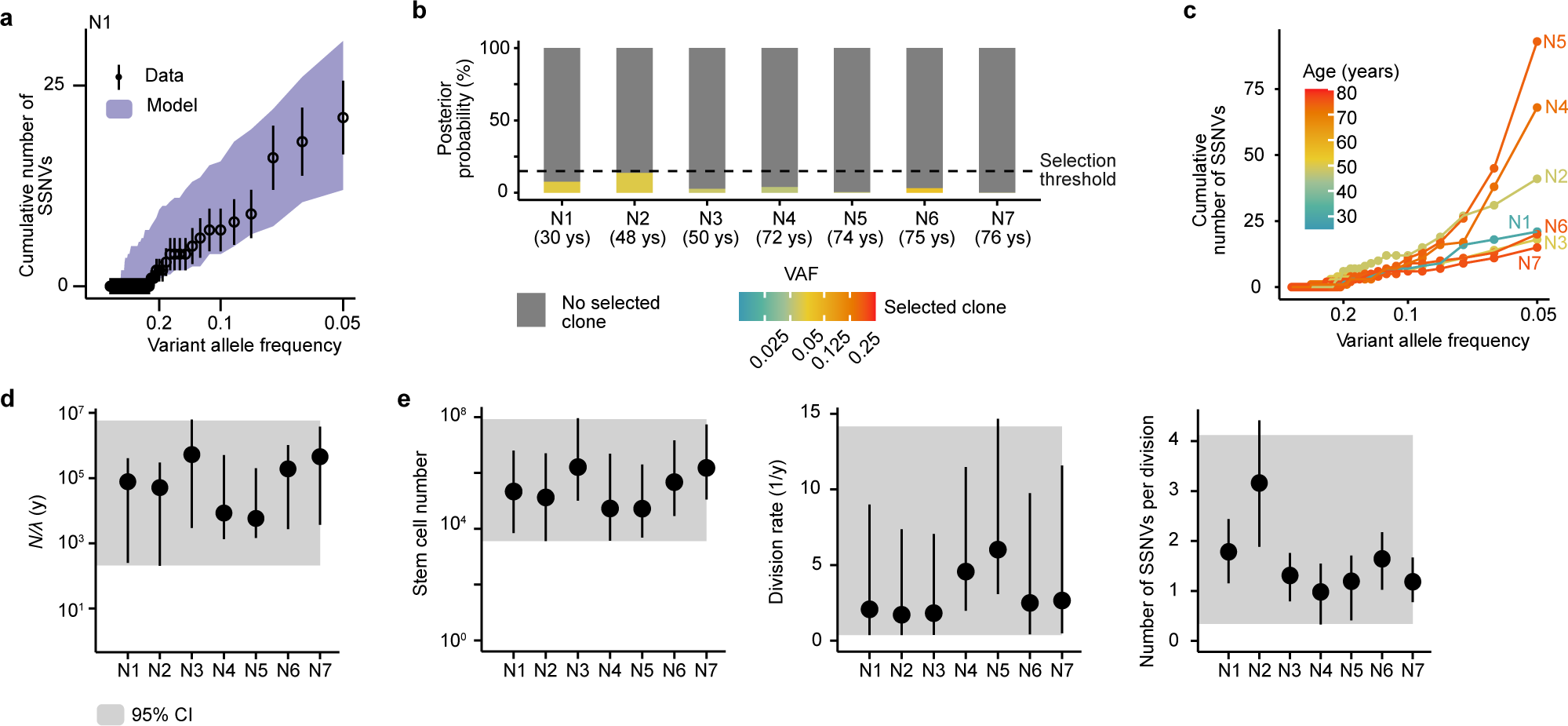
Neutral evolution in physiological hematopoiesis measured by bulk WGS. **a**, Model fit to the cumulative 1/VAF distribution measured in DNA from CD34^+^ HSPCs of sample N1 (points and error bars, mean and standard deviation of the measured data; purple area, 95% posterior probabilities of the model fit). **b**, Model support for clonal selection (posterior probabilities conditioned on clones with VAF ≥5%) and neutral evolution across seven samples without known CH driver mutation (N1-N7). The color scale shows the inferred VAF of selected clone. The dotted lines shows the 15% selection threshold. All cases are classified as neutrally evolving. **c**, Cumulative VAF distributions for samples N1-N7 shown in **b**. Age of individuals from whom samples N1-N7 were obtained are color coded. **d**, Inferred ratio between stem cell number and division rate (*N/λ*). **e**, Inferred parameter estimates for the number of hematopoietic stem cells (left panel), division rate of a stem cell per year (middle panel) and the number of new SSNVs acquired per cell division for samples N1-N7 shown in **a**. Shown are median and 80% credible intervals for each sample; the grey areas give the 95% confidence interval across the lower and upper bounds of all samples.

In samples N1-N3 and N6-N7 there were around 20-30 SSNVs with VAF ≥5% whilst N4 and N5 stood out with >60 SSNVs (Fig. 4c). To understand this, we computed the ratio of HSC number to HSC division rate, *N*/*A* (Fig. 4d). This parameter characterizes the timescale over which neutral evolution will cause a variant to reach fixation. For all our neutral samples, except N4 and N5, SCIFER inferred *N*/*A* to be ∼10^5^ to 10^6^ years, supporting the notion that clones do not drift to large size in normal human hematopoiesis^24^. However, individuals N4 and N5 had smaller *N*/*A*, of ∼10^4^ years, consistent with genetic drift increasing SSNV counts at low VAF (cf. Fig. 1e-f). To probe the differences in HSC behavior between N4/N5 and the other individuals in more detail, we inferred stem cell number (5x10^4^-1.5x10^6^ HSCs), HSC division rate (1-2 times a year for N1-N3 and N6-N7) and rate of acquired SSNV/cell division (1-3) (Fig. 4e). Interestingly, these estimates were broadly consistent with our inferences from the pseudo-bulk data from published samples (Fig. 3d-f), although the credible intervals were substantially larger. However, N4 and N5, were above 70 years of age (Fig. 4b), showed larger HSC division rates than the other individuals (Fig 4e), about 5 self-renewing divisions per year, suggesting that the drift effects seen at low VAF could be caused by more rapid HSC proliferation.

In summary, the analysis of the individuals with neutral HSC dynamics supports the notion that large hematopoietic clones do not arise by drift. However, in two of the seven cases we inferred acceleration of HSC division, by unknown causes. This likely accounts for increased cumulative number of SSNVs at lower VAFs, which we interpret as a signature of clonal drift.

### Clonal selection frequently linked with accelerated stem cell self-renewal

Next, we quantified selection for the 10 individuals that had at least one mutation in either *DNMT3A*, or *TET2*, or *ASXL1*, genes (2 males, 8 females, ranging between 62 and 89 years of age; Supplementary Table 2) and where SCIFER identified clonal selection (Fig. 5a-b; Extended Data Fig. 7a) consistent with the known CH driver mutation. The clone size inferred by SCIFER, without knowledge of the driver, agreed with the VAF of the driver (Fig. 5c). We identified no CNVs or SVs associated with CH (Extended Data Fig. 6a; Supplementary Tables 4 and 5). The single base-pair variant profiles were consistent with those observed in patients without CH driver mutations, indicating that the same mutational processes were active (Extended Data Fig. 6b).

**Fig. 5.**
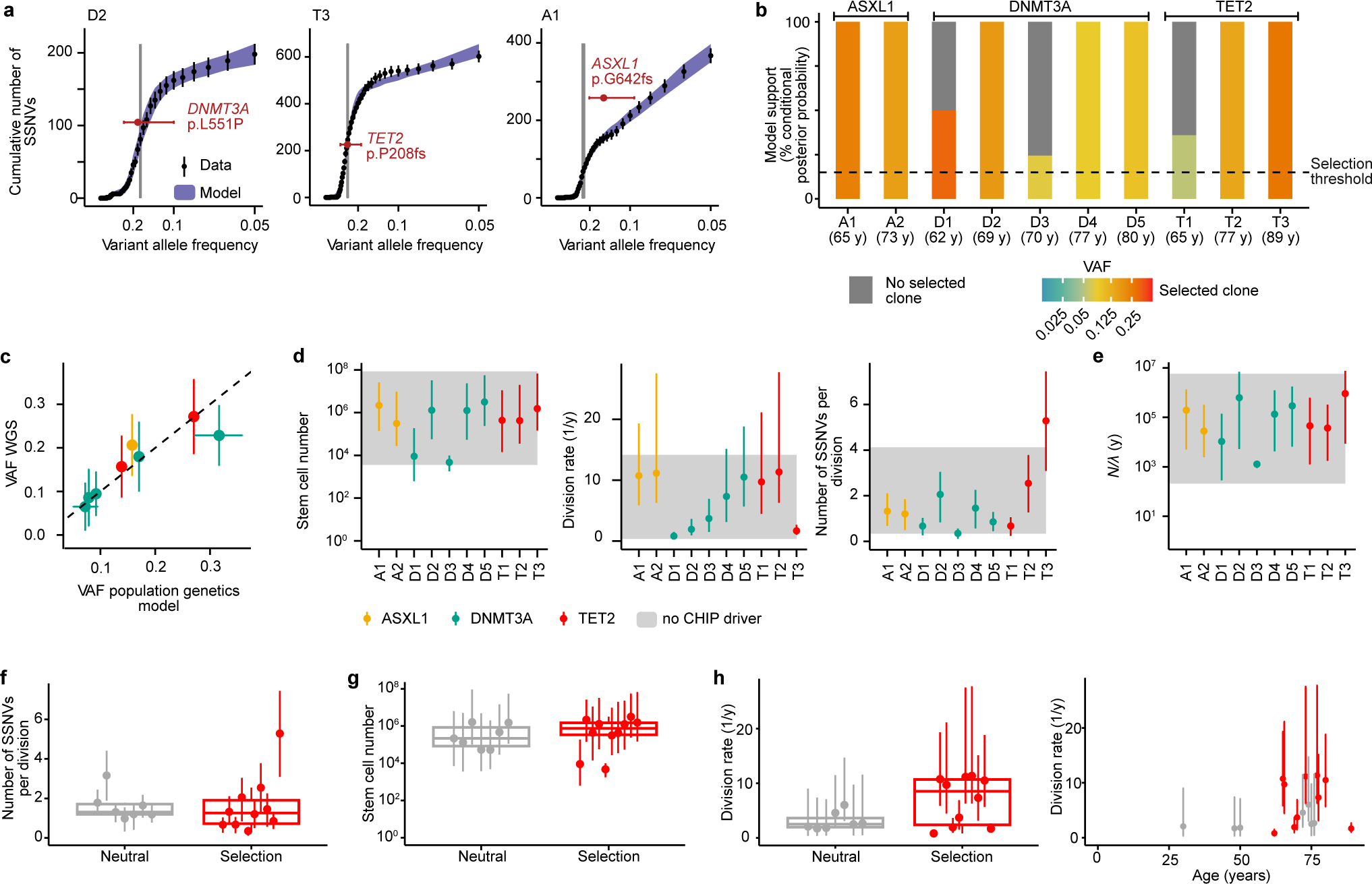
Clonal selection in hematopoiesis measured by bulk WGS. **a**, Model fit to the cumulative VAF distribution measured in CD34^+^ HSPCs of samples D2, T3 and A1 (points and error bars, mean and counting errors of the measured data; purple area, 95% posterior probabilities of the model fit; grey area, 80% credible interval of the estimated clone size; red points and error bars, mean and 95% confidence interval of the VAF of known CH drivers). **b**, Model support for clonal selection (posterior probability conditioned on clones >5% VAF) and neutral evolution in ten cases classified as selected and with a CH driver mutation in *AXSL1* (A), *DNMT3A* (D) and *TET2* (T). The age of the individuals is shown below each bar; the color scale shows the inferred VAF of the selected clone and the dotted lines shows the 15% selection threshold. **c**, Estimated sizes of the selected clones (median and 80% credible intervals) versus measured VAF of the known CH driver (mean and 95% CI according to a Binomial distribution). **d**, Estimated number of HSCs contributing to hematopoiesis (left), their division rate (middle) and number of SSNVs acquired per cell division (right) for the ten cases shown in **b**. Shown are median and 80% credible intervals for each sample; the grey areas give the 95% confidence interval across the lower and upper bounds of the seven neutrally evolving samples without known CH driver for comparison (c.f. Fig. 4). **e**, As in **d**, but showing the ratio between stem cell number and division rate (*N*). **f**, Estimated number of newly acquired SSNVs per HSC division in the 7 neutrally evolving cases (introduced in Fig. 4) as compared to 10 cases with selection for a known CH driver (introduced in this figure; points and error bars, median and 80% credible intervals for each sample; boxplots, median and interquartile range, whiskers extend to the largest and smallest value no further than 1.5 times the interquartile range). **g**, Estimated number of HSCs contributing to hematopoiesis in the 7 neutrally evolving cases (introduced in Fig. 4) as compared to 10 cases with selection for a known CH driver (introduced in this figure; points and error bars, median and 80% credible intervals for each sample; boxplots, median and interquartile range, whiskers extend to the largest and smallest value no further than 1.5 times the interquartile range). **h**, Estimated HSC division rate in the 7 neutrally evolving cases (introduced in Fig. 4) as compared to 10 cases with selection for a known CH driver (introduced in this figure; Points and error bars, median and 80% credible intervals for each sample; boxplots, median and interquartile range, whiskers extend to the largest and smallest value no further than 1.5 times the interquartile range). The right panel plots the estimated division rate against the age of the individuals from whom samples were obtained. Samples that are neutrally evolving are shown in grey, samples with evidence of clonal selection are shown in red.

We asked whether HSC parameters differed between cases with distinct driver genes. There was no association between the CH driver gene and the total number of HSCs (Fig. 5d, left panel), the HSC division rate (Fig. 5d, middle panel) and the rate of acquisition of new SSNV per cell division (Fig. 5d, right panel). Two samples with a *DNMT3A* mutation (D2 and D4) stood out with a smaller HSC number (∼5x10^3^) and one of them, D4, also had a lower *N*/*1* ratio (Fig. 5e). Otherwise, the HSC number estimate was rather uniform, whereas the HSC division rate showed considerable variation.

Next, we asked whether there were systematic differences compared to HSC parameters in individuals with neutral evolution (N1-N7; Fig.4d,e). The rate of acquisition of new SSNV per cell division and HSC numbers were overall not distinguishable between individuals who showed neutral evolution versus those with CH (Fig. 5f,g). By contrast, the HSC division rates were more variable in individuals with CH. The majority of samples with selected clones (7 out of 10) had a higher HSC division rate than the neutral-evolution group, while three CH cases had HSC division rates comparable to cases of neutral evolution (Fig. 5h, left panel).

When plotting HSC division rate as a function of age, we noticed that larger division rates were concentrated in the older individuals (≥65 years of age). The two individuals exhibiting neutral evolution with a higher HSC division rate also fell into this older age group (N4 and N5, cf. Fig. 3e). However, the largest division rates (≥10 divisions per year) were exclusively found in cases where selection operates. Hence, our data suggest that variability in stem cell division rate increases with age, and that clonal selection events are frequently linked with accelerated HSC division.

### Clonal selection without known drivers

Finally, we analysed the five cases (U1-U5) where SCIFER identified major selected clones that did not have a corresponding driver mutation at the same VAF in the panel of 97 known CH-associated genes. In all five samples, copy number profiles (Extended Data Fig. 8a) and single base pair substitution profiles (Extended Data Fig. 8b) were as for all the other samples. For U1, U3, U4 and U5, SCIFER found clear evidence of selection (Fig. 6a-b). By contrast, the model support for selection for sample U2 was just above the threshold for selection; it should be kept in mind that the false-positive rate is ∼20% for small clones at the detection threshold of 5% VAF (Fig. 2a).

**Fig. 6.**
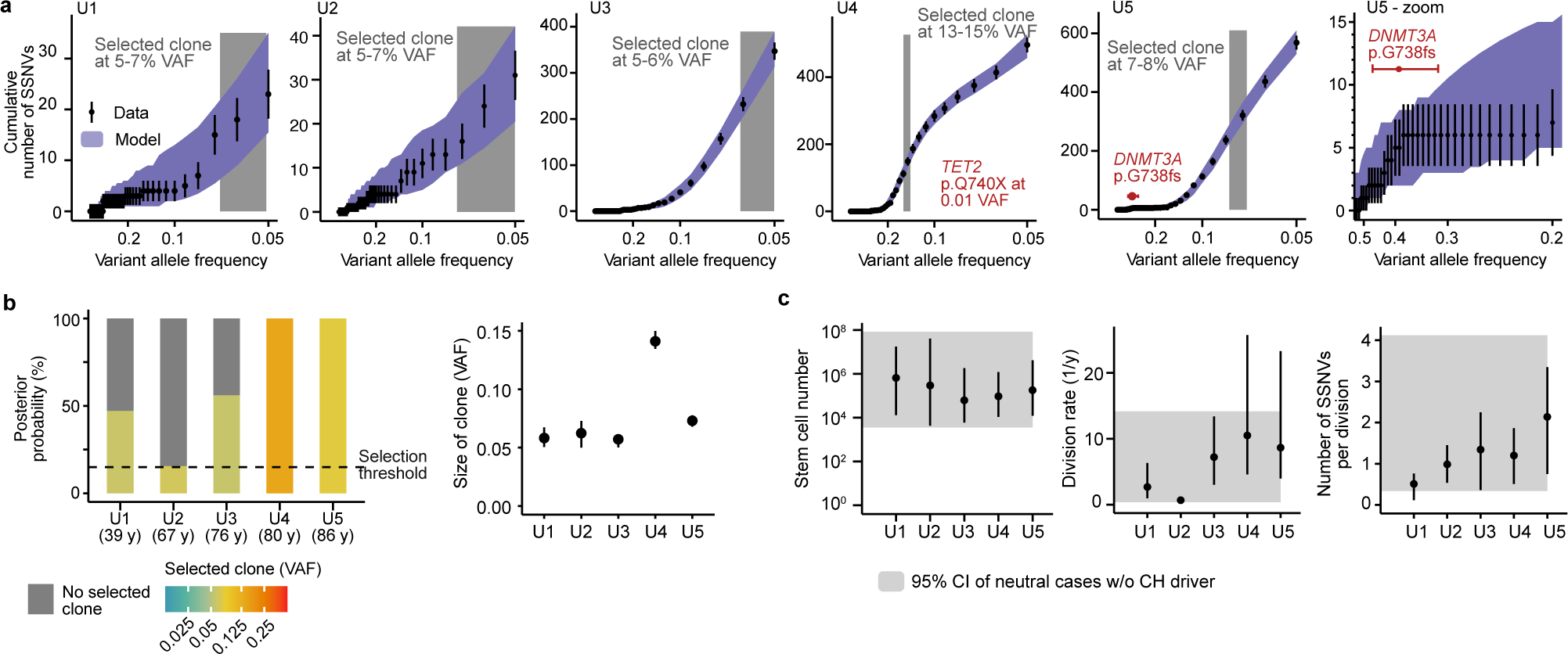
Clonal selection for unknown drivers. **a**, Model fits to the cumulative 1/VAF distribution measured in CD34^+^ HSPCs of samples U1-U5 (points and error bars, mean and standard deviation of the measured data; purple area, 95% posterior probabilities of the model fit; grey area, 80% credible interval of the estimated clone size). In U4 and U5, red points and error bars show mean and 95% confidence interval of the VAF of known CH driver mutations (*TET2* for U4 and *DNMT3A* for U5). In both U4 and U5 selection was not associated with the known CH driver mutations (U4, inferred selected clone at 14% VAF and *TET2* mutation at 1% VAF; U5, inferred selected clone at 7% VAF and *DNMT3A* mutation at 43% VAF). The panel on the right shows an expanded view of the cumulative VAF distribution and the model fitting to the data at high VAF. At most 6 SSNVs were acquired prior to the DNMT3A mutation, suggesting that the mutation was acquired early during development. **b**, Model support for clonal selection (posterior probability conditioned on selected clones with ≥5% VAF) and neutral evolution across samples U1-U5. The age of the individuals is shown below each bar; the color scale shows the inferred VAF of selected clone and the dotted lines shows the 15% selection threshold. The right panel gives the estimated clone size (median and 80% credible intervals). **c**, Estimated number of HSC contributing to hematopoiesis stem cell number (left), HSC division rate (central) and number of newly SSNVs acquired per cell division (right). Shown are median and 80% credible intervals for each sample; for comparison, the grey areas give the 95% confidence interval across the lower and upper bounds of neutrally evolving samples without known CH driver (as shown in Fig. 4).

Samples for U1-U3 did not have CH mutations on panel sequencing. Samples for U4 and U5 had recurrent mutations in known CH driver genes (*TET2* for U4 and *DNMT3A* for U5). However, the size of the leading clone inferred by SCIFER differed strongly from the VAF of the respective CH driver. In U4 the SCIFER-inferred clone (14% VAF) was much larger than the VAF of the *TET2* mutation (1%). Hence, a large HSC clone had already grown before the *TET2* mutation occurred. In U5, the *DNMT3A* mutation was acquired very early in life, most likely during embryogenesis as there were only 6 clonal SSNVs (Fig. 6a, amplified right panel) and almost reached fixation (43% VAF). Within this early selected clone, SCIFER inferred a subclone with a VAF of 7.5% without a known CH driver. This interesting result indicates that SCIFER may miss selection events occurring in embryogenesis (such as the *DNMT3A* mutation in U5) because of the small clonal SSNV count (note that a clone of similar size, but much later origin and hence larger SSNV count, was readily detected by SCIFER in the pseudo-bulk sample id2259^22^; Extended Data Fig. 2g).

Assessing the HSC parameters (Fig. 6c), we found little variation in HSC number (Fig. 6c, left panel) and overall similar numbers as in the CH cases with known drivers (Fig. 5d). However, three individuals (U3-U5) had accelerated HSC divisions, with a rate close to 10 divisions per year, while the other two cases had a rate similar to the majority of non-CH cases (about 2 divisions per year). The accelerated rate for U3-U5 agreed with the HSC division rate of the majority of CH cases with known drivers (Fig. 5h). These data further support the link between clonal selection events, with known or unknown drivers, and accelerated HSC turnover.

### Homogeneous occurrence of selection events across life

Finally, we asked when clones with or without known CH drivers originated in life and how fast they expand. The clone ages inferred by SCIFER showed that clones of both kinds typically emerged several decades before BM sampling (Fig. 7a); moreover, there were no significant differences in clone age between associated driver mutations in *DNMT3A, TET2* and *ASXL1* and unknown drivers (Fig. 7b). The time-averaged clonal growth rates showed substantial inter-individual variation, even within the five individuals with a *DNMT3A* mutation and the three individuals with a *TET2* mutation (Fig. 7c). We did not detect systematic differences in growth rates for clones associated with different drivers or unknown drivers (Fig. 7d).

**Figure 7.**
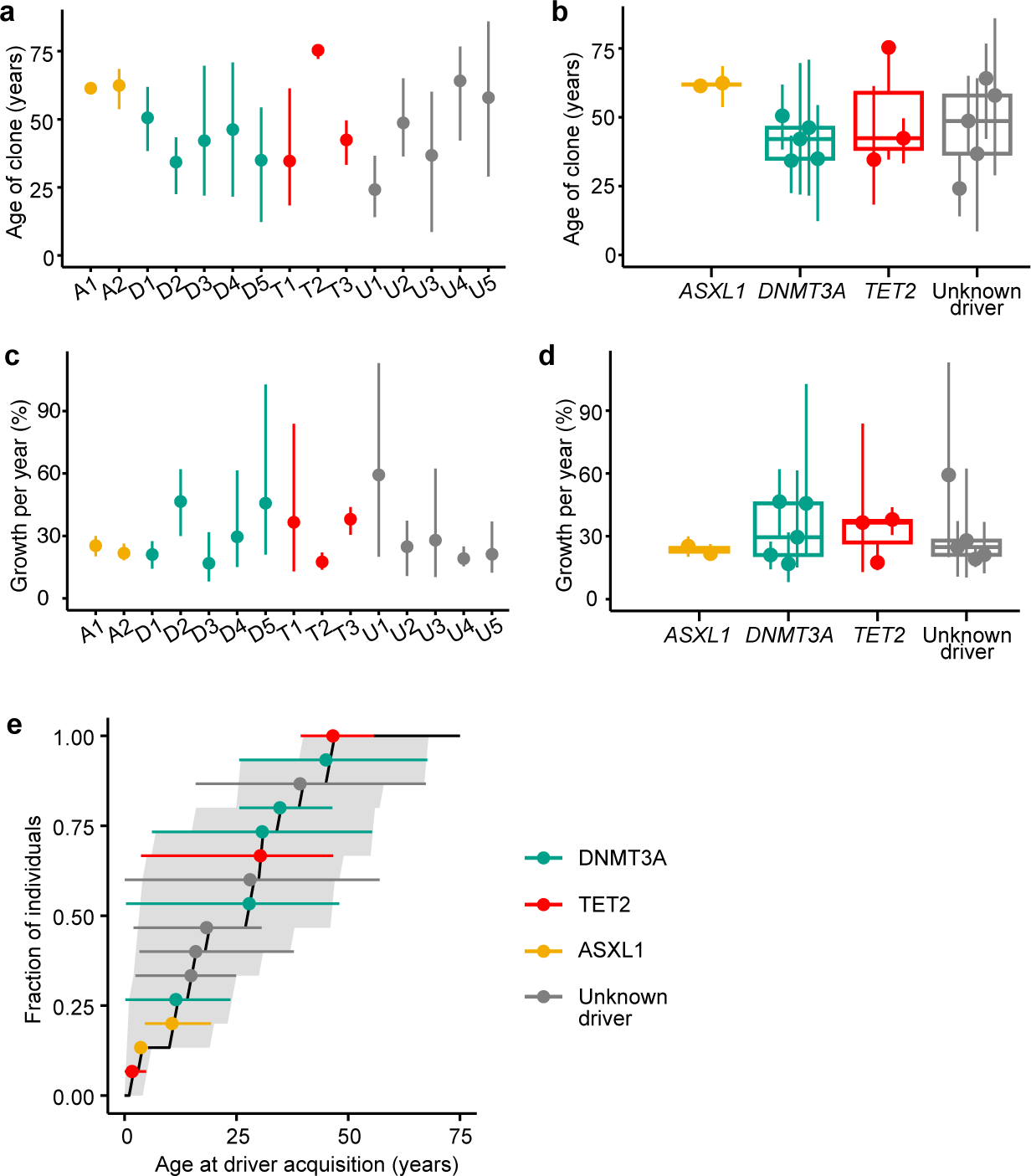
Selection dynamics with and without known drivers. **a**, Estimated age of the selected clone (for the ten cases with known drivers introduced in Fig. 5 and the five cases with unknown drivers introduced in Fig. 6; points, median; error bars, 80% credible intervals). **b**, Per-driver-gene comparison of the estimated age of the selected clone; data are the same as in **a** (points, median; error bars, 80% credible intervals; boxplots, median and interquartile range, whiskers extend to the largest and smallest value no further than 1.5 times the interquartile range). **c**, Estimated growth rates for the fifteen cases introduced in **a** (points, median; error bars, 80% credible intervals). **d**, Per-driver-gene comparison of the estimated growth rates for the fifteen cases introduced **a** (points, median; error bars, 80% credible intervals; boxplots, median and interquartile range, whiskers extend to the largest and smallest value no further than 1.5 times the interquartile range). **e**, Cumulative distribution of estimated age at CH driver acquisition (median); shaded area, lower and upper bounds of the cumulative distribution of estimated age at CH driver acquisition based on 80% credible intervals of the estimated parameters; points and error bars, median and 80% credible intervals for the per-sample estimates. Data are from ten cases with selection of known CH driver mutations and five cases with selection of unknown drivers introduced in **a**.

A key question is when driver mutations of CH arise in life; previous work based on phylogenetic reconstruction has inferred that expanded clones originate before 40 years of age ^26^. Combining all our cases with clonal selection, which were selected in our cohort without reference to the age of the individual, we deduced the age-incidence distribution for the onset of clonal selection (Fig. 7e). Intriguingly, we found that selected clones were born at a fairly constant rate from early childhood to about 50 years of age. This limit does not imply that selection events no longer occur beyond this age; it may rather be a limit of clone detection (e.g., a clone that grows by 25% per year requires 40 years to grow to detectable size, ∼0.05x10^5^ cells, for 90x bulk WGS; the oldest individual in our cohort is 89 years of age, and hence such a clone born at age 49 would just be detectable). Hence our data suggest that selection events in HSCs occur quite uniformly across life.

## Discussion

Selection of mutant somatic cells is central to cancer evolution. In contrast, neutral evolution was initially emphasized in non-malignant tissues such as the normal gut epithelium by quantitative studies of stem cell dynamics^10^. However, more recent data show that, over the long human lifespan, selection also shapes somatic mosaicism in non-malignant tissues, and this can be accompanied by altered tissue function^22,24-26,35^. Here, we present a method to detect clonal selection and quantify the dynamic parameters of tissue stem cells from snapshot WGS data using a single bulk tissue sample. Our population-genetics approach, applied to populations of cells, provides a quantitative framework to model homeostatic competition of clones arising during development or adult homeostasis. Akin to recent quantitative approaches to cancer evolution^29-31,36-38^, our theory delineates how the allele frequency spectra of somatic variants are shaped by neutral evolution versus selection. In contrast to previous work on distinguishing genetic drift from selection in expanding tumors ^30,36^, our study of somatic evolution in homeostatic tissues requires the explicit treatment of genetic drift and its interplay with clonal selection. This allows us to identify selection in an individual homeostatic tissue without knowledge of the underlying driver events.

An alternative for detecting selection in a normal tissue agnostic of the driver is provided by phylogenetic reconstruction from the genomes of individual stem and progenitor cells^2^. With ∼300 HSPCs per individual analyzed, this method has yielded detailed insight into clonal structure of hematopoiesis in 12 individuals, thus far^22,26^. These studies have highlighted multiple selected clones in an individual (each selected clone corresponding to an expanded clade in the phylogeny and the smallest expanded clade having a frequency of ∼1.3%). The majority of expanded clones had no known driver events. While less sensitive at sequencing depths of 90x we used here, our method is cost-effective, and hence scalable, enabling a search for unknown drivers in larger population cohorts. Moreover, if we sequence at 270x our simulations suggest SCIFER would achieve a similar level of sensitivity as phylogenetic reconstruction. SCIFER can also be applied to tissues where stem-cell clones cannot be grown easily from stem cells.

Moreover, the agreement of basic quantitative parameters of hematopoiesis – HSC number, turnover and mutation rate – inferred with the help of the phylogenetic tree^2^ and from SCIFER is remarkable, as our method strongly relies on variants with higher VAF emerging relatively early, whereas the phylodynamic inference strongly relies on the large number of low-VAF SSNVs generated later. This suggests that these HSC parameters are rather uniform across time, at least before selection events occur.

Our study is the most comprehensive to date, of quantitative parameters of human HSC dynamics in humans. We find neutrally evolving hematopoiesis in individuals of all ages up to our clonal detection limit. In individuals without CH, we find uniform dynamics of HSCs across life, except for two older individuals (72 and 74 years, respectively) with smaller numbers of stem cells. For these two individuals, we infer noticeable drift of variants, generated in fetal development and early in life, to allele frequencies detectable in bulk WGS. This observation cautions against solely using the number of SSNVs per se, as evidence for selection^27^. In several individuals with selected clones, HSC division and mutation rates are as in the cases of neutral evolution, indicating that CH can arise from normal HSC dynamics.

Remarkably, however, we found an enrichment for cases with increased time-averaged HSC division rate of HSC in individuals with CH. We also found a uniform rate of acquiring CH across life. This includes CH driver events occurring early in life or even in embryogenesis (such as the *DNMT3A* mutation in case U5), akin to blood cancer (myeloproliferative disorder^39^).There are at least three avenues of future research that arise from this observation. First, the accelerated HSC division may aid the emergence of CH by increasing the rate of driver mutation acquisition^21^. Second, as SCIFER estimates an average division rate for all HSC in an individual, it leaves open the question of whether the HSC division rate is different between HSCs that are selected, compared to those that are not selected, within a CH marrow. Recent single cell data suggest that all HSCs within in a CH bone marrow environment have transcriptional signatures of inflammation but that this is less pronounced in selected HSCs^33^. This raises the hypothesis that selected HSCs with a dampened response to inflammation, may function better in an inflammatory environment^40^. Third, as our data are time averaged from single timepoint measurements it would be fascinating to apply SCIFER suggest to measure HSC parameters (division rates and birth of clones) serially in longitudinal samples across an individual’s lifetime. Specifically, it would be important to test the impact of specific stressful events such as intercurrent inflammation, infection and disease and therapeutic interventions. Given the scalability of SCIFER, this is now feasible.

Consistent with previous studies^22,25,26^ we identified five cases with clonal selection in the absence of known drivers. The selective advantage conferred by unknown driver events was indistinguishable from known CH drivers, and unknown drivers were also frequently associated with increased HSC self-renewal rate. Although in-depth analyses of the genome-wide mutation profiles revealed potential candidates for driver mutations in four of these cases (Supplementary Table 6), the statistical power of the current study is not large enough to identify new recurrent driver mutations with confidence. However, by applying SCIFER to large cohorts it may be possible to more accurately determine the frequency of individuals with unknown driver events, where selection is operative in blood or other tissues. With this information in hand, the phenotype and clinical significance of these events could be assessed. Combined with in depth genome-wide multi-ome data and functional screens, this may also provide insight into the mechanisms by which these currently unknown driver events mediate selection and shape tissue function.

**Extended Data Fig. 1.**
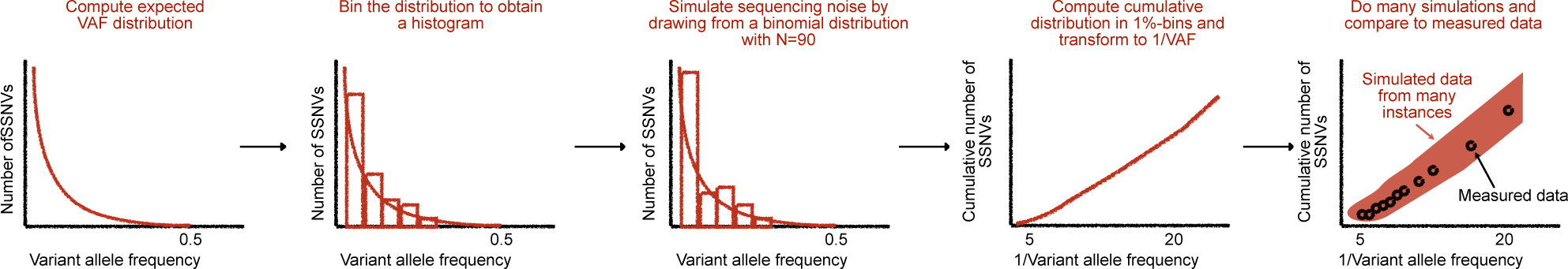
Quantifying selection and drift from bulk whole genome sequencing data. Parameter estimation with approximate Bayesian computation. First, the expected variant allele frequency histogram is analytically computed. Then, sequencing noise is simulated by drawing from a binomial distribution with average 90x coverage. The modelled cumulative distribution is compared to the measured data for varying VAFs (1% step size).

**Extended Data Fig. 2.**
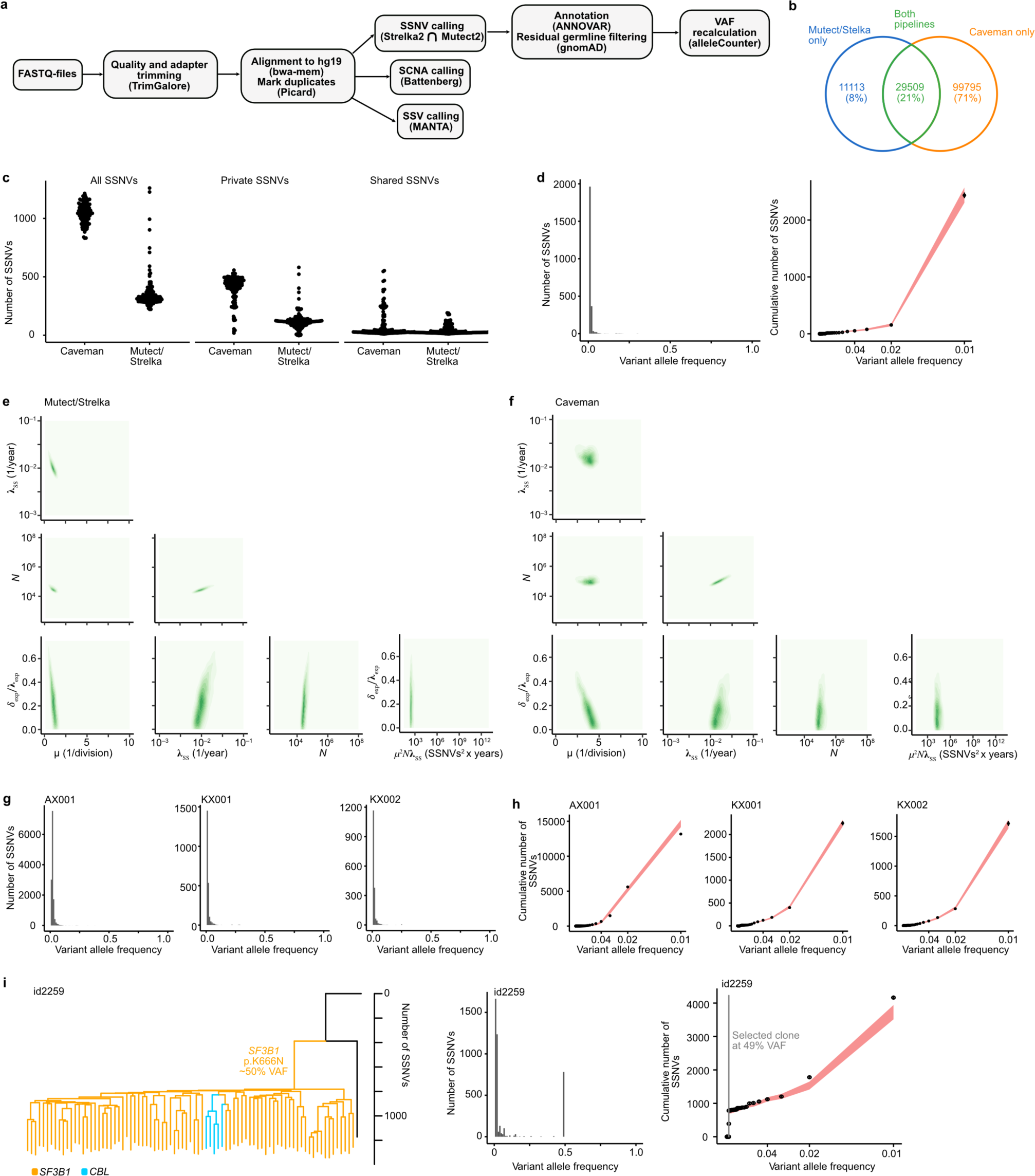
Quantifying drift and selection in published WGS data from single HSPC clones. **a**, Somatic variant calling pipeline (using hg19). SSNV, somatic single nucleotide variant. SCNA, somatic copy number abnormality. SSV, somatic structural variant. **b**, Number of shared and unique SSNVs identified in 140 whole genomes from HSPC clones from published data^2^ identified by Caveman (original publication) and our pipeline (intersection of Mutect and Strelka shown in **a**). **c**, Number of SSNVs of different classes identified by each variant caller. **d**, Left panel, variant allele frequency distribution (truncated at 0.01 VAF) of SSNVs identified by Caveman from published data^2^. Right panel, model fit to the cumulative 1/VAF distribution shown in left panel (points and error bars, mean and standard deviation of the measured data; red area, 95% posterior probabilities of the model fit). **e**, Two-dimensional posterior probability distributions for the parameters estimated by SCIFER from the cumulative 1/VAF distribution of SSNVs identified with our pipeline (intersection of Mutect and Strelka) in published data^2^. Axis limits show the range of the prior distributions. **f**, Two-dimensional posterior probability distributions for the parameters estimated by SCIFER from the cumulative 1/VAF distribution of SSNVs identified with Caveman in published data^2^. Axis limits show the range of the prior distributions. Note that while the posterior for the fraction of cell loss, *δ*_exp_′λ_exp_, is relatively broad, this has only a small effect on the values of the mutation rate and also the other parameters (see Supplementary Note 1). **g**, VAF distribution (truncated at 0.01 VAF) of SSNVs from 1150 genomes from three samples (AX001, KX001 and KX002) that did not show signs of clonal selection (SSNVs taken without re-call from published data^26^). **h**, Model fit to the cumulative VAF distribution shown in **g** (points and error bars, mean and standard deviation of the measured data; red area, 95% posterior probabilities of the model fit). **i**, Left, phylogenetic tree of published sample id2259, containing a large selected clone with a *SF3B1* pK605N mutation (and a smaller *CBL* subclone)^22^ (reconstruction from the original publication). Middle panel, VAF distribution (truncated at 0.01 VAF) of SSNVs (taken without re-call from^22^). Right panel, model fit to the cumulative 1/VAF distribution (points and error bars, mean and standard deviation of the measured data; red area, 95% posterior probabilities of the model fit; grey area, 80% credible interval of the estimated clone size).

**Extended Data Fig. 3.**
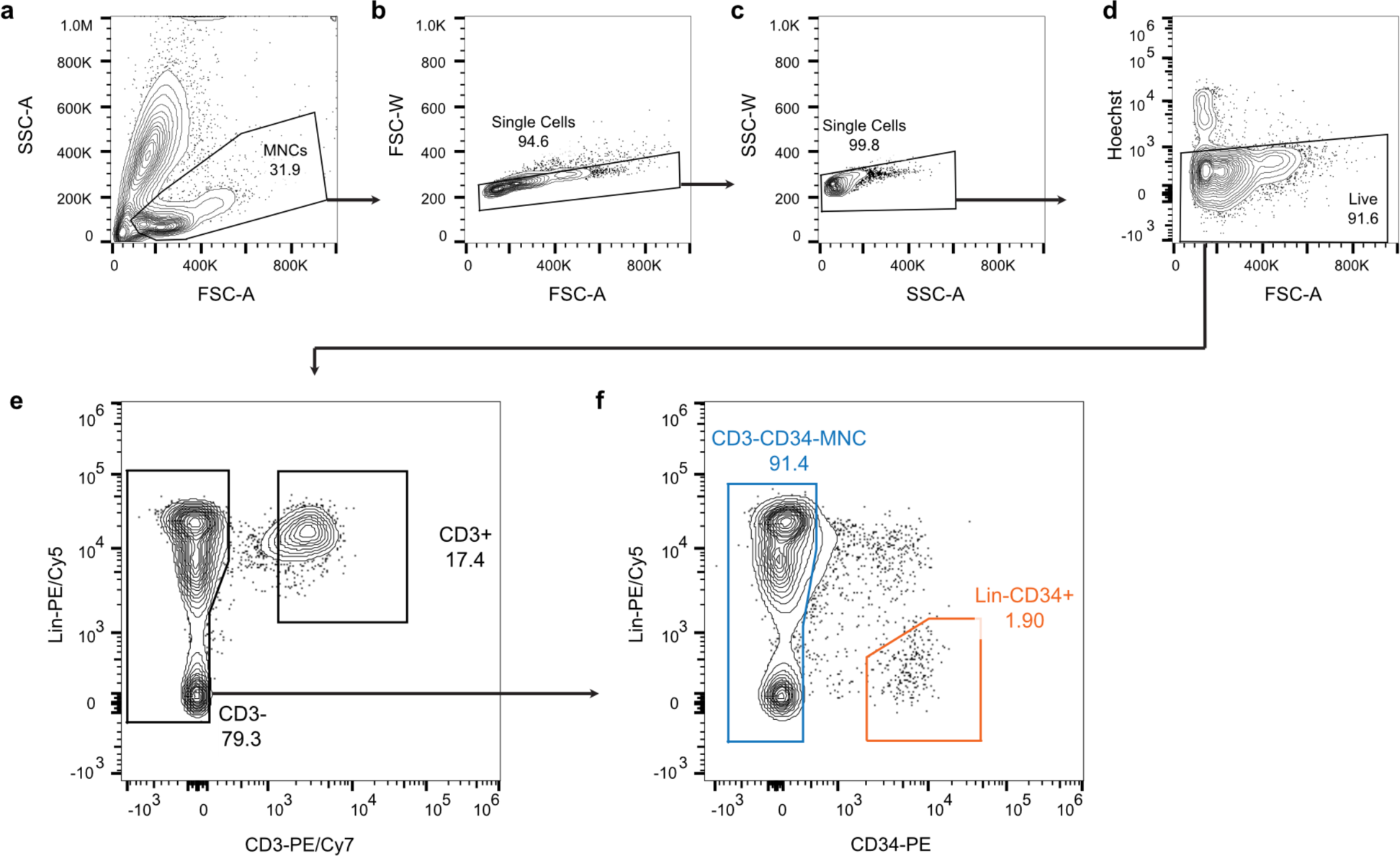
FACS gating for hematopoietic stem and progenitor cells. **a-f**, Cells were stained with a panel of antibodies (Methods), then single and live mononuclear cells were sorted (MNCs, **a-d**). T cell depleted MNCs (MNC(–T)) were sorted from the CD3-, CD34+ cell fraction (**e** and **f**), and, hematopoietic stem and progenitor cells were sorted from the CD3-Lin-CD34+ cell fraction (**f**). Image created with FlowJo v10.8.1; values in **a**-**f** give percentages.

**Extended Data Fig. 4.**
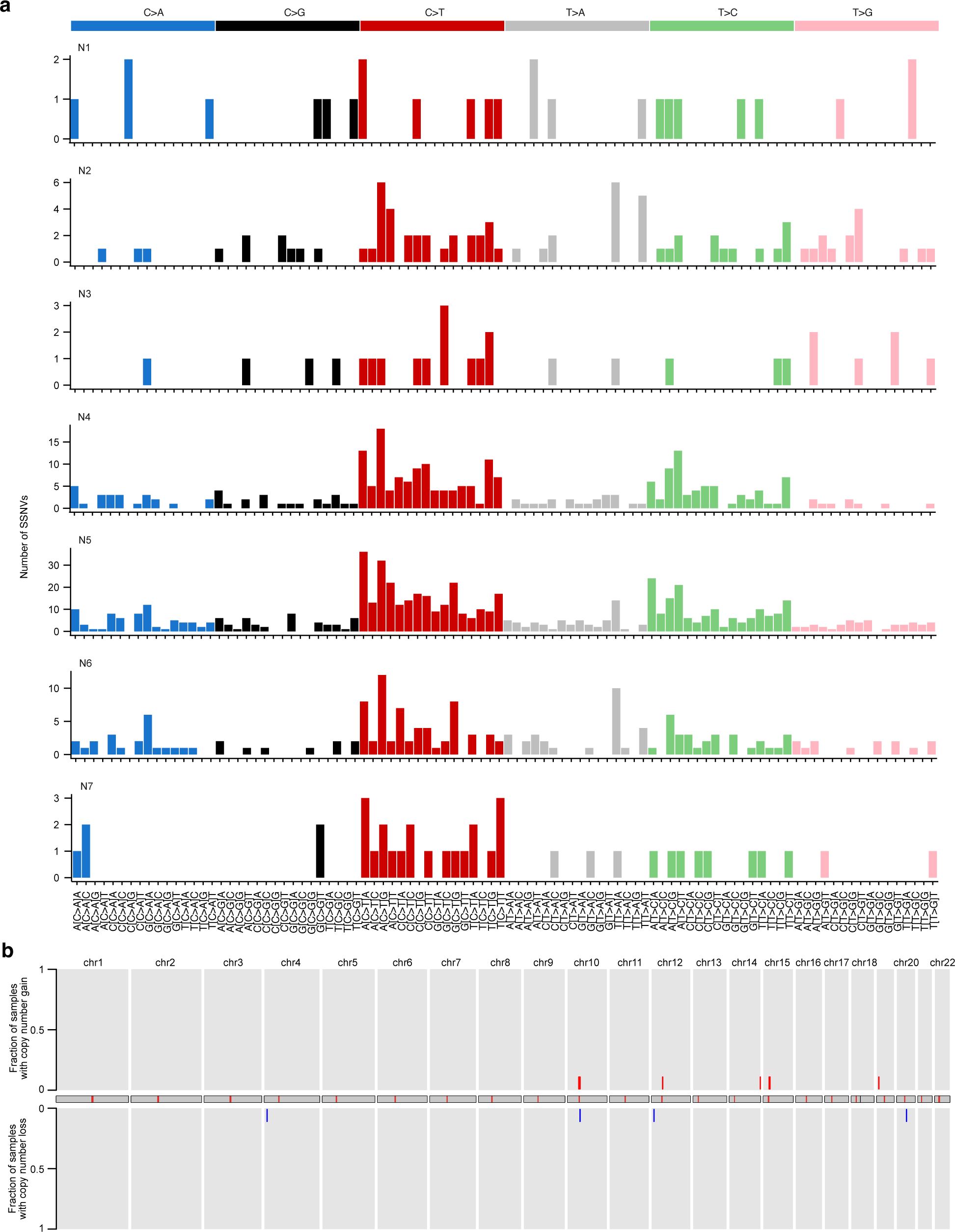
Somatic variants in individuals with neutral evolution. **a**, Trinucleotide substitution patterns across the genome in CD34^+^ HSPC in samples N1-N7. **b**, Genome-wide (chromosomes indicated and arrayed across x-axis) profiles of copy number gains (red) and losses (blue) in CD34^+^ HSPC across the 7 same samples as in **a**.

**Extended Data Fig. 5.**
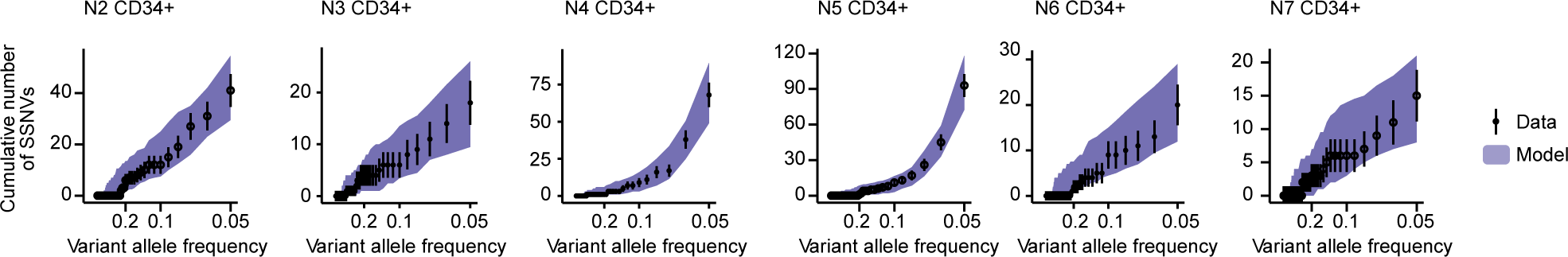
Quantifying neutral drift and mutation accumulation from bulk whole genome sequencing. Model fits to the cumulative VAF distributions measured by whole genome sequencing (WGS) of CD34^+^ HSPC samples from six individuals (N2-N7) without known CH driver mutation (points and error bars, mean and counting errors of the measured data; purple areas show the 95% posterior probabilities of the model fit). The model fits show no evidence of selection.

**Extended Data Fig. 6.**
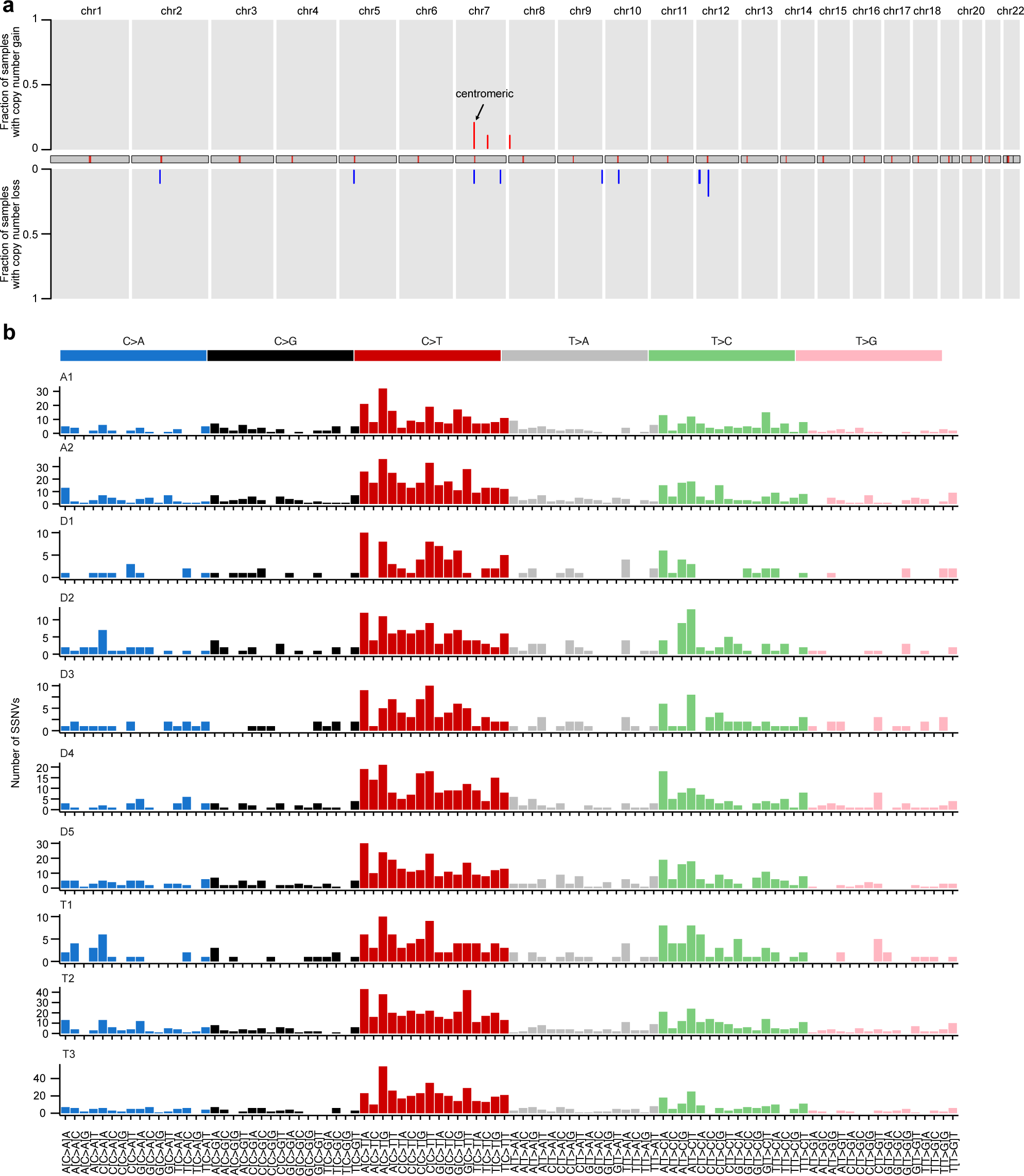
Somatic mutations in clonal hematopoiesis. **a**, Genome-wide (chromosomes indicated and arrayed across x-axis) profiles of copy number gains (red) and losses (blue) in CD34^+^ HSPC from 10 individuals (A1-A2, D1-D5 and T1-3) with a known CH driver mutation. **b**, Genome wide trinucleotide substitution patterns in CD34^+^ HSPC across the 10 same samples as in **a**.

**Extended Data Fig. 7.**
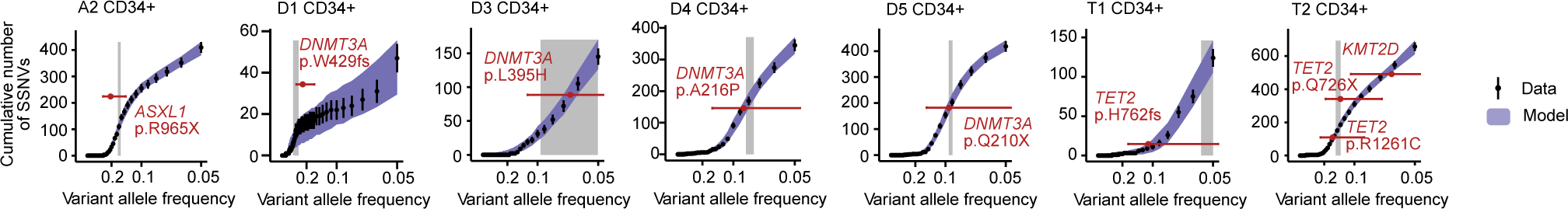
Quantifying clonal selection from bulk whole genome sequencing. The cumulative VAF distributions measured by whole genome sequencing (WGS) of CD34^+^ HSPC from seven individuals with known CH driver mutations and evidence for clonal selection (A2, D1, D3-5 and T2-3; A=individual with *ASXL1* mutation; D=individual with *DNMT3A* mutation; T=individual with *TET2* mutation) (points and error bars, mean and counting errors of the measured data). The violet areas show the 95% posterior probabilities of the model fit. Grey areas, 80% credible interval for the estimated clone size; red points and error bars, mean and 95% confidence interval of the VAF of known CH drivers

**Extended Data Fig. 8.**
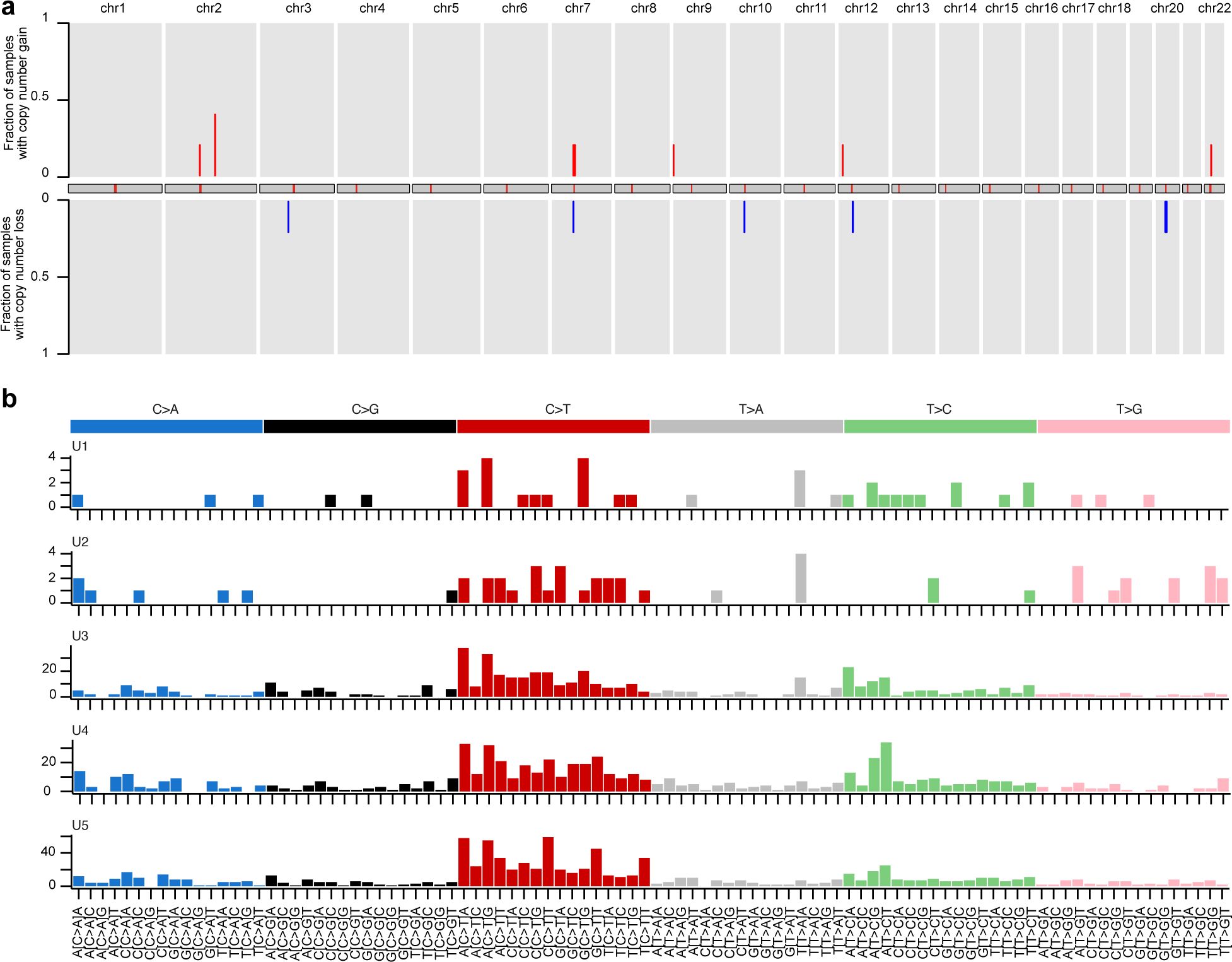
Somatic mutations in clonal hematopoiesis with selection of unknown drivers. **a**, Genome-wide (chromosomes indicated and arrayed across x-axis) profiles of copy number gains (red) and losses (blue) in CD34^+^ HSPC from 5 individuals (U1-5) where expanded selected clones were detected but no known genetic driver was identified. **b**, Genome wide trinucleotide substitution patterns in CD34^+^ HSPC across the 5 same samples as in **a**.

## Methods

### Ethical approval

Patient samples were collected with informed consent under the Mechanisms of Age-Related Clonal Haematopoiesis (MARCH) Study. Written informed consent was obtained from all participants in accordance with the Declaration of Helsinki. This study was approved by the Yorkshire & The Humber -Bradford Leeds Research Ethics Committee (REC Ref: 17/YH/0382).

### Study samples

Participants were recruited from individuals undergoing elective total hip replacement surgery at the Nuffield Orthopaedic Centre, Oxford. Exclusion criteria were: history of rheumatoid arthritis or other inflammatory arthritis, history of septic arthritis in the limb undergoing surgery, history of hematological cancer, bisphosphonate use, and oral steroid use. Patient characteristics are summarized in Supplementary Table 2. At the time of surgery, trabecular bone fragments and BM aspirates were obtained from the femoral canal and collected in anticoagulated buffer containing acid-citrate-dextrose, heparin sodium and DNase. Samples of peripheral blood were collected in EDTA vacutainers. Hair follicle samples were collected from participants as a germline control. Peripheral blood and BM mononuclear cells (MNCs) were isolated by Ficoll density gradient centrifugation and viably frozen. Peripheral blood granulocytic cell pellets were frozen for later DNA extraction. Genomic DNA was extracted from bone marrow MNCs, peripheral blood granulocytes and hair follicles using a DNeasy Blood & Tissue Kit (Qiagen).

### Cell sorting

Thawing media was prepared with IMDM medium (Gibco) supplemented with 20% fetal bovine serum (FBS) and 110 µg/mL DNase. BM samples were thawed at 37°C in a water bath, 1 mL warm FBS was added, and the suspension then diluted by dropwise addition of 8 mL thawing media. The suspension was centrifuged at 400 g for 10 mins, cells were resuspended in flow cytometry staining medium (IMDM with 10% FBS and 10 μg/mL DNase), filtered through a 35 μm cell strainer, and placed on ice.

Cells were stained with the following antibodies: anti-CD34-PE (1:160, Biolegend, clone 581), anti-CD3-PE/Cy7 (1:100, Biolegend, clone HIT3a), anti-CD2-PE/Cy5 (1:160, Biolegend, clone RPA-2.10), anti-CD4-PE/Cy5 (1:160, Biolegend, clone RPA-T4), anti-CD7-PE/Cy5 (1:160, Biolegend, clone CD7-6B7), anti-CD8a-PE/Cy5 (1:320, Biolegend, clone RPA-T8), anti-CD11b-PE/Cy5 (1:160, Biolegend, clone ICRF44), anti-CD14-PE/Cy5 (1:160, eBioscience, clone 61D3), anti-CD19-PE/Cy5 (1:160, Biolegend, clone HIB19), anti-CD20-PE/Cy5 (1:160, Biolegend, clone 2H7), anti-CD56-PE/Cy5 (1:80, Biolegend, clone MEM188), and anti-CD235ab-PE/Cy5 (1:320, Biolegend, clone HIR2). Following antibody incubations, cells were washed with 1 mL flow cytometry staining buffer, centrifuged at 350 g for 5 min and resuspended in flow cytometry staining buffer containing 1:10,000 Hoechst 33342 live/dead stain.

Cell sorting was performed on a BD FACSAria Fusion or Sony MA900 equipped with a 100 µm nozzle or sorting chip. Unstained, single-stained, and FMO controls were used to determine background staining and compensation in each channel. Doublets and dead cells were excluded. The following populations were sorted with a mean purity > 95%: Lin^−^CD34^+^ HSPCs, Lin^+^CD3^+^ T cells, and Lin^+/–^CD34^−^CD3^−^ MNCs. Genomic DNA from sorted cell populations was extracted using a QIAamp DNA Micro Kit (Qiagen).

### Whole genome sequencing

#### Library preparation and sequencing

Sequencing libraries were prepared using the Illumina DNA Prep Kit (#20018704) and Nextera DNA CD Indexes (#20018707) according to manufacturer’s guidelines. The Qubit HS DNA Assay Kit (Invitogen, #Q32854) and a tape station run with the Agilent HS D5000 Assay Kit (#5067-5593) were used for quality control. Thereafter, libraries were diluted to 10nM, pooled and sequenced on either HiSeq X PE150 or on NovaSeq 6000 PE150.

Low quality bases on raw sequencing reads were trimmed using Trim Galore v0.5.0 (https://www.bioinformatics.babraham.ac.uk/projects/trim_galore/) with cutadapt v2.8^41^ with the following settings: –quality 30, –illumina –length 32 –trim-n –clir_R1 2 –clip_R2 2 – three_prime_clip_R1 2 –three_prime_clip_R2 2. Trimmed read pairs were mapped to the human reference genome (build 37 with the GRCh37.75 genome annotation) using bwa mem v0.7.12^42^. Mapped reads were coordinate-sorted with samtools sort v1.5^42^, followed by marking duplicate read pairs with gatk MarkDuplicates v4.0.9.0^43^ and indexing with samtools index.

#### Detection of SSNVs and indels

Somatic SNVs and small insertions/deletions in BM and blood samples were called with Strelka v2.9.2^44^ and Mutect2, GATK v4.2.0.0^43^, using the matched hair follicle samples as germline control. Variants in repeat regions and simple repeat regions (downloaded from UCSC table browser, setting the assembly to hg19, the track to “RepeatMasker” or “Simple Repeats”; accession date: December 5^th^, 2018) were filtered using bedtools intersect v2.24.0^45^. To generate a set of high-confidence variants, only variants that passed the default filters of both Strelka and Mutect2 were retained. To identify remaining germline variants, variants were looked up in dbSNP (v150) and population frequency of reported variants was annotated based on the Genome Aggregation Database (gnomAD v2.1.1; https://gnomad.broadinstitute.org/). Variants were then filtered to exclude potential contamination of germline variants based on the global allele frequency (AF) of gnomAD. Specifically, only variants with AF <0.01 were retained as somatic variants. In a few cases, a CH driver was identified by panel sequencing at low VAF, but was not recovered by Strelka and Mutect; here, we re-examined the respective position using bcftools mpileup v1.10.2^46^. All variants were annotated with ANNOVAR (version May2018; http://annovar.openbioinformatics.org/)^47^ according to the human genome build version 19. For SSNVs, VAFs were re-calculated directly from the BAM files using alleleCounter v.4.0.2 (https://github.com/cancerit/alleleCount; with default base and mapping quality thresholds) and MaC (https://github.com/nansari-pour/MaC).

#### Detection of copy number variants

Genome-wide subclonal copy number aberrations (CNA) were identified with Battenberg (v2.2.10; https://github.com/Wedge-lab/battenberg), which has been described in detail previously^48^.

#### Detection of structural variants

Structural variants (SVs) from whole genome data were identified using MANTA v1.6.0^49^ with default tumor-normal pair settings where samples were compared to the germline control to identify somatic structural rearrangements. We kept variants that passed Manta’s default filter, had a minimal SOMATICSCORE of 30, were not classified as IMPRECISE, had at most 2 variant reads in the control sample, and had VAF <5% in the control sample.

#### Driver analysis

To comprehensively search for candidate driver mutations, we concatenated published lists of putative driver genes in clonal hematopoiesis and leukemia from

- The cancer driver database Intogen^50^ (subsetting genes described in “ALL”, “AML”, “CLL”, “CML” or “MM” from release 2020.02.01, and the clonal hematopoiesis gene list from Pich et al., 2022^51^),
- The cancer driver database Cosmic^52^ v94 (subsetting on genes with annotation in leukemia),
- Bick et al., 2019^53^ and Jaiswal et al., 2014^12^,
- A curated list of clonal hematopoiesis-associated mutations compiled from five large studies^12,13,17,54,55^.

Mutations identified by Strelka and targeting any of these genes were looked up in ClinVar^56^ (https://ftp.ncbi.nlm.nih.gov/pub/clinvar/vcf, downloaded on March 31^st^, 2023) and annotated accordingly. Using the R package drawProteins^57^ v1.18.0 we then collected information on the protein domain targeted by each variant. Moreover, we manually annotated variant information (involvement in disease, functional evidence for mutations at this site, SNPID, association with genetic discorders and whether the variant targets a functional domain) using the manually curated variant information “homo_sapiens_variation.txt” from Uniprot^58^ (downloaded on April 4^th^, 2023), variant information provided on www.uniprot.org, and variant information on www.omim.org^59^. Thereafter, we computed SIFT^60^ and Revel^61^ prediction scores for each substitution. We kept variants

- causing a “stopgain”, “stoploss”, “frameshift_insertion”, or “frameshift_deletion”, or
- with “Conflicting interpretations of pathogenicity”, “Likely pathogenic”, “Pathogenic/Likely pathogenic” or of “uncertain significance” according to ClinVar, or
- annotated to a related disease (based on uniprot.org or omim.org) or
- annotated as pathogenic (uniprot.org) or
- targeting a “binding site” or a “DNA binding” site (uniprot.org) or
- with a REVEL score ≥0.75 or
- targeting either *ASXL1*, or *DNMT3A* or *TET2*.

This yielded a set of variant positions, based on which we recalled variants in all blood and control samples with bcftools mpileup^46^ v 1.10.2, using the option -p, and bcftools call, using the option -mA. In the final set, we kept variants targeting any of *ASXL1, DNMT3A* or *TET2* and variants that were found on at least 2 reads in the sample and on <5 reads in the controls.

### Analysis of single-cell WGS data

We tested our population genetics model on published single-cell WGS data from refs.^2,22,26^. To this end, we computed VAFs from the single-cell phylogenies as

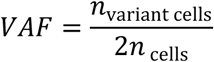

where *n*_variant cells_ is the number of sequenced cells harboring a variant *n*_cells_ is the total number of sequenced cells; the factor 2 accounts for diploidy.

#### Parameter estimation

We fit our population genetics model to the cumulative VAF distribution truncated at 0.01, as detailed below and using the prior probabilities outlined in Supplementary Table 7.

#### Re-analysis of SSNV and indels

To exclude potential differences between our data and the published data that may arise due to differences in the variant calling pipelines, we re-called SSNVs and indels in the data from ref.^2^. In particular, we used the intersection between Mutect2^43^ and Strelka2^44^, filtered remaining germline variants using gnomAD (retaining variants with AF <0.01) and re-calculated VAFs directly from the BAM files using alleleCounter v.4.0.2 (https://github.com/cancerit/alleleCount; with default base and mapping quality thresholds) and MaC (https://github.com/nansari-pour/MaC). We filtered the remaining variants as stated in ref.^2^. In brief, we removed variants occurring in >120 of the 140 colonies, variants that fell within 10bp of each other and variants with a coverage <6 on autosomes or <3 on sex chromosomes in >5 samples. Moreover, we retained only variants with a mean VAF > 0.3 across all samples with at least one mutant read. Finally, we excluded sites at which >10% of the samples with at least one mutant read had a VAF<0.1.

#### Phylogenetic reconstruction

We reconstructed single-cell phylogenies based on the re-called SSNVs and Indels from Lee-Six et al.^2^ by converting the mutation table into a fasta file, learning the phylogenetic tree using MPBoot v1.1.0^62^ and plotting the tree using custom scripts from^26^ (https://github.com/emily-mitchell/normal_haematopoiesis/).

### Population genetics model

#### i Theory

We modeled the evolution of variant allele frequencies mechanistically, accounting for accumulation, drift and selection of somatic variants in a homeostatic tissue. The model is parametrized by the rate at which HSCs divide during adulthood, λ, the number of SSNVs acquired between two cell divisions, μ, the number of HSCs during adulthood, *N*_ss_, as well as the time of origin of the selected clone, *t*_s_, and its selective advantage expressed by *r*. We bundled the model functions in an R package, SCIFER, available on https://github.com/VerenaK90/SCIFER/tree/paper.

##### Modeling the site frequency spectrum of somatic variants generated by neutral evolution

The time-dependent site frequency spectrum *S*.(*t*) gives the number of variants with clone size *i*, where *i* ranges from some minimally observable clone size (owing to WGS sequencing depth) up to the total number of stem cells *N*(*t*). We derive analytical expressions for the site frequency spectra resulting from neutral evolution and clonal selection and compare these with measured VAF histograms.

To begin with neutral evolution, we develop a stochastic model for accumulation and drift of neutral somatic variants during developmental expansion and subsequent homeostasis of the stem cell pool. The model assumes that stem cells proliferate via symmetric self-renewing divisions, with rate λ, and are lost by differentiation and cell death, with rate *δ*.Between two subsequent stem cell divisions, on average μ new variants are introduced into each daughter cell. These variants are inherited to daughter cells, and, depending on the dynamics of the corresponding stem cell clone, may either go extinct, or drift to variable frequencies. The site frequency spectrum generated by these dynamics is

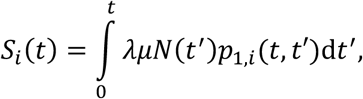

describing the generation of new variants at time *t*′, in a population of size *N*(*t*′), and their drift to clone size *i* up to the time of measurement *t* with probability *p*_1,._(*t*, t′). To compare *S*_*i*_(*t*) with measured VAF histograms, we transform from clone size to VAF:

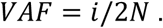

We will see below that the transformed site frequency spectrum depends on the parameters of the birth-death process and mutation acquisition as follows:

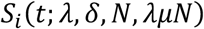

Proliferation and loss rate, as well as the total cell number, appear as individual parameters algebraically and can in principle be inferred from the data as individual parameters. Hence also the mutation rate, appearing in the product λμ*N*, can be inferred. Practical identifiability will depend on the sequencing depth (Fig. 2c). Thus our approach will aim to qualitatively distinguish between neutral evolution and selection in experimental data and determine the underlying parameters quantitatively.

To describe developmental expansion of the stem cell pool followed by homeostasis, we concatenated linear birth-death processes. During development, the division rate will exceed the loss rate, λ_exp_ > *δ*_exp_, hence defining a supercritical birth-death process. At time *t*_1_, the system reaches its steady-state with a constant number of active stem cells, *N*_ss_. The cellular dynamics are now appropriately described by a critical birth-death process with steady-state rate λ_ss_= *δ*_ss_.

To compute the site frequency spectrum during developmental expansion, we consider the probability that a cell acquiring a new variant will expand to a clone of size *a* within time *t*:^63^

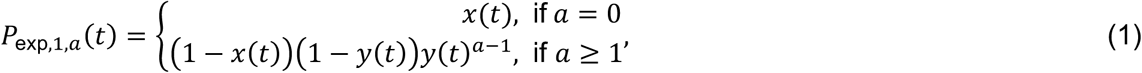

With

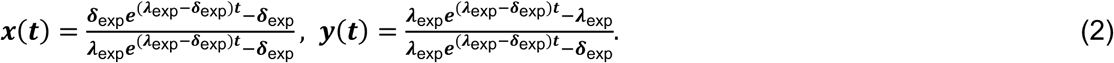

The measured VAF histograms report on the number of variants with a given frequency. To calculate variant number in the model, we note that between time *t*′ and *t*′ + d*t*′ on average μ λ_exp_*N*(*t*′)d*t*′ variants are generated. Hence,

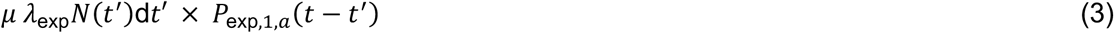

variants introduced at time *t*′ each occur in *a* cells at time *t*.

Next, we consider the fate of these variants during tissue homeostasis. Then stem cell division and loss will be balanced, occurring both at steady-state rate λ_ss_. Accordingly, the drift of already existing variants is described by the critical birth-death process. Recall that *t*_1_ is the time at which the homeostatic stem cell number is reached. A variant in *a* cells at *t*_1_ will drift to occur in *b* cells within time *t* with probability^63^

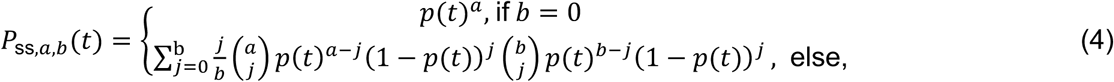

With

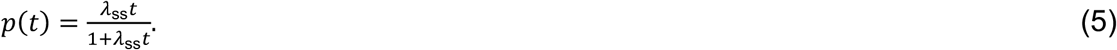

Thus, at *t* ≥ t_1_, the number of variants occurring exactly in *b* cells is

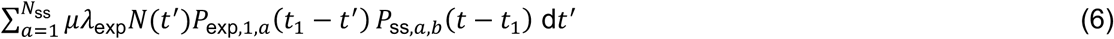

for variants generated during developmental expansion (*t*′ < t_1_).

Finally, we consider variants acquired during homeostasis, which evolve according to the critical birth-death process entirely. Hence, the number of such variants occurring exactly in *b* cells is

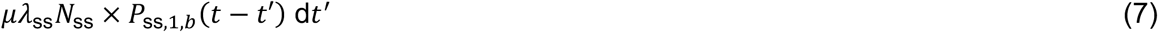

where *t*′ ≥ t_1_. Combining the contribution of variants originating in expansion or homeostasis, we arrive at the site frequency spectrum of neutral variants in a homeostatic tissue without selection:

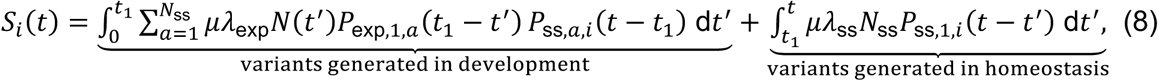

Eq. (8) generalizes a result by Ohtsuki and Innan^64^ for expanding tissues.

##### Modeling the site frequency spectrum of somatic variants under selection

Clonal selection will modify the site frequency spectrum. Consider that a positively selected mutation is acquired at time *t*_s_ and reduces the rate of stem cell loss by the factor *r* < 1 (the alternative for imparting selective advantage, an increase in the rate of stem cell division, yields very similar results). The cell number of mutated stem cells, *C*(*t*), expands at the expense of the number of normal stem cells, *U*(*t*) = N_ss_ − *C*(*t*), according to the competition model

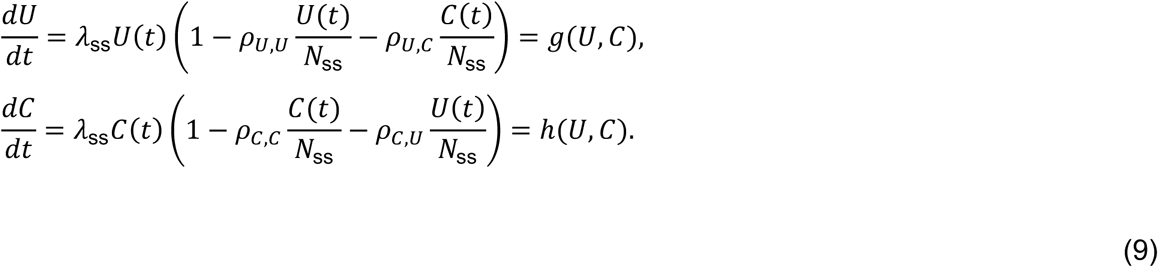

The *ρ* parameters denote phenomenological competition coefficients between and within the mutant clone and the normal stem cells, maintaining homeostasis. We have no further growth if either normal or mutated stem cells fill the entire compartment, *g*(*N*_ss_, 0) = 0 = *h*(0, N_ss_) and *g*(0, N_ss_) = 0 = *h*(*N*_ss_, 0), implying that *ρ*_*U,U*_ = 1 = *ρ*_*C,C*_. Moreover, *ρ*_*C,U*_ = r and hence

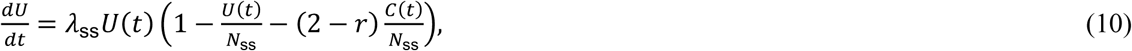

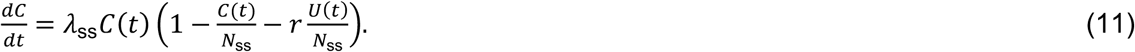

The number of mutant stem cells *C*(*t*) will remain much smaller than the number of normal stem cells for extended periods of time, and hence we approximate Eq. (11) by

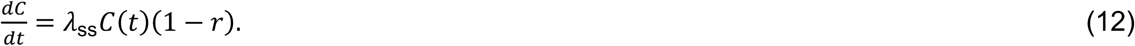

Eqs (10) and (12) will be used when computing the site frequency spectrum.

With clonal selection, the site frequency spectrum has three principal contributions for somatic variants originating (i) before the positively selected mutation occurred, *S*_*i*,1_, (ii) after this mutation occurred and happening in the mutant clone, *S*_*i*,2_, and (iii) after the mutation occurred but happening in normal stem cells, *S*_*i*,3_:

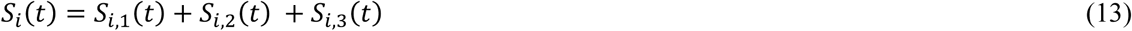

We specify each contribution in turn. The site frequency spectrum of variants in the mutant clone, *S*_.,2_, is

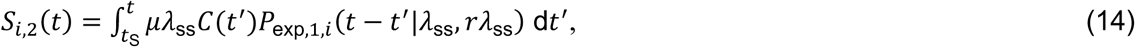

where *P*_exp,1,._ is given by Eq. (1).

The site frequency spectra of variants acquired in normal stem cells, *S*_.*i*,1_ and *S*_.*i*,3_, are shaped by the decline of the number of normal stem cells after the emergence of the selected clone at *t*_S_ (Eq. 10). We approximate this process by a subcritical birth-death process with division rate λ_ss_ and effective loss rate *δ*_eff_(*t*). The loss rate is chosen such that the expectation of the decline of normal stem cells in the birth-death process equals the number of normal stem cells lost by the competition dynamics, *D*. (Eq. 10) implies that

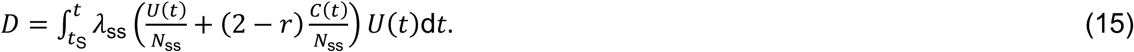

Therefore, the effective loss rate *δ*_eff_(*t*) is defined by

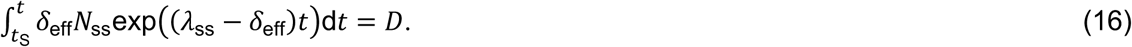

The site frequency spectrum of variants acquired in normal stem cells after *t*_S_, *S*_.*i*,3_, is the superposition of (*N*_ss_ − 1) independent linear subcritical birth-death processes and is given by

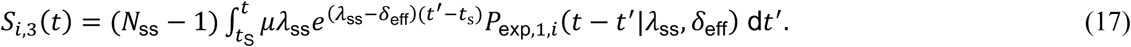

Finally, to compute the site frequency spectrum of variants acquired prior to the generation time of the selected mutation *t*_S_, *S*_.*i*,1_, we denote by *k* the size of the clone harbouring a particular variant at *t*_*S*_. With probability *k/N*_ss_, the founder cell of the mutant clone will have this variant (as well as *k* − 1 normal stem cells at *t*_S_); with probability 1 − *k/N*_ss_, the variant will not be in the founder cells of the mutant clone (as well as *k* normal stem cells will harbor it at *t*_S_). Noticing that clones of size *i* < *C*(*t*) can only contain normal stem cells, we find that

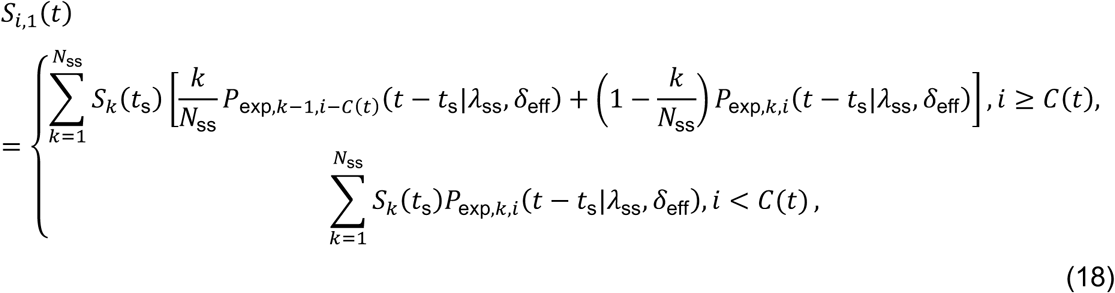

where *S*_*k*_(*t*_*S*_) is the site frequency spectrum at *t*_*S*_ and *P*_exp,*a,b*_ is the clone size distribution generated by a subcritical birth-death process initiated by *a* cells:^63^

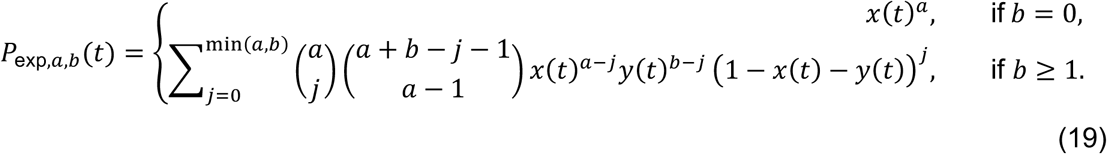

Thus, we have specified the contributions to the site frequency spectrum of a stem cell population containing a selected clone, *S*.(*t*) = S_.,1_(*t*) + S_.,2_(*t*) + S_.,3_(*t*).

#### ii Parameter estimation

Parameter estimates were obtained for each donor separately, using approximate Bayesian computation as implemented in pyABC^65^ (posterior sample size of 1000 and termination criterion *ε*_min_ *=* 0.15). We performed the following steps to compare the cumulative number of variants, 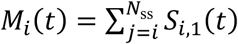, between measured and simulated data:

1. [Experimental data] Determine *M*_*i*_(*t*) for the experimental data. For bulk WGS data, include variants if they are supported by at least 3 reads, absent in the germline control sample, and if the locus is covered by at least 10 and at most 300 reads. For pseudo-bulk WGS data, include all variants. Denoting the measured variant allele frequencies with VAF, compute the cumulative number of variants, *M*_experimental,*f*_ = ∑ *n*_VAF*≥f*_, where *n*_*j*_ denotes the number of variants with VAF given by *j*. We use bins for *f* of width 0.01, with 1.15 ≤ *f* ≤ 1 for bulk WGS data (30x and 90x), and 1.11 ≤ *f* ≤ 1 for pseudo-bulk WGS data and bulk WGS data (270x, simulated).
2. [Model] Sample parameters from their prior distributions (Supplementary Table 7).
3. [Model] Simulate the expected cumulative number of variants, *M*_sim,*F*_(*t*_*a*_), where *t*_*a*_ is the age of the patient (for numerical implementation of the model see **Section iv** below). The bins are as with the experimental data. Depending on the following criterion, based on the prior parameter sample, the cumulative VAF histogram is simulated with either the neutral model or the selection model: If a selected clone could grow above the detection limit with the prior parameter sample, the selection model is used; otherwise the neutral model is used. Formally, if 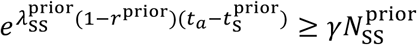 then the selection model is used. The detection limit depends on the sequencing depth, with γ = 0.15 for 90x and 30x WGS data, and γ = 0.11 for pseudo-bulk WGS data and simulated 270x bulk WGS data. Smaller clones are not accessible by the respective sequencing approach and can hence be neglected. Importantly, note that the selection model can return a posterior corresponding to neutral evolution.
4. [Model] Add the number of variants present in the founder cell of the hematopoietic system, 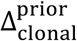, to *M*_sim,1_ (corresponding to mutations acquired during early development).
5. [Model] Simulate experimental error of sequencing. To this end, generate the expected site frequency spectrum, *S*_sim_(*f*_*i*_) = *M*_sim_(*f*_*i*_) − *M*_sim_(*f*_*i*+1_). For bulk WGS data, sample for each simulated variant a sequencing coverage *c* according to *Pois*(ĉ), *ĉ* ∈ {30, 90,270}, depending on the sequencing depth. Thereafter, sample VAFs for each variant according to 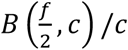, where *B* denotes the Binomial distribution. Discard variants supported by less than 3 reads. For pseudo-bulk WGS data, sample VAFs for each variant according to *B*(*f n*_*cells*_)/(2*n*_*cells*_), where *n*_*cells*_ is the number of sequenced single-cell clones. Compute the sampled cumulative number of variants *M*_sim,*F*,sampled_.
6. [Model vs Experimental data] Determine the distance function for ABC.

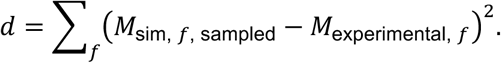

#### iii Classification of cases as neutrally evolving or as selected

We classified cases as selected if 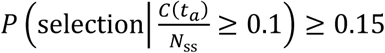, that is if the selected clone has at least 5% VAF and the corresponding posterior probability was at least 15%. These thresholds were defined based on in silico generated test data, and correspond to the WGS depth of 90x (Figure 2). Upon sample classification, we computed the 80% highest density intervals for each parameter on the parameter subsets supporting neutral evolution or clonal selection, respectively.

#### iv Numerical computation of the cumulative number of variant allele frequencies

To facilitate numerical computation of the cumulative number of variants, *M*_*i*_(*t*), in particular if *N* becomes large, we implemented numerical approximations that are detailed in the following.

##### Drift of somatic variants acquired during expansion

First, we assessed the site frequency spectrum after exponential expansion, *M*_*i*,exp_(*t*_1_). Recall that *t*_1_ denotes the time point of transition between expansion and homeostasis. Accordingly,

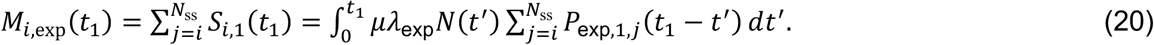

As *N*_ss_ is large, we approximate the sum in Eq. (20) by integration, yielding

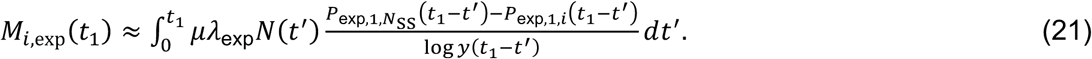

The full site frequency spectrum is obtained by evaluating Eq. (21) for every clone size *i* with 1 ≤ *i* ≤ *N*_ss_. However, for large *N*_ss_, this becomes numerically costly (the upper bound of the prior for *N*_ss_ is 10^8^). Hence, we evaluate 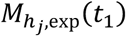 according to Eq. (21) for a series of bins with logarithmic bin sizes for small clones (<10%), and constant bin sizes of 5% for clones > 10%, as follows:

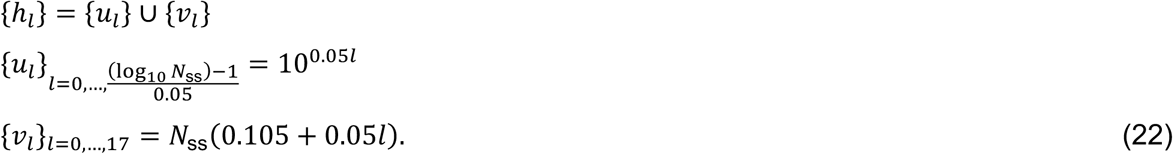

We determined a binned site frequency spectrum, 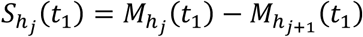, at the end of the expansion phase. Then the cumulative site frequency spectrum of variants acquired during expansion at a later time point *t*_2_ > t_1_, is obtained as:

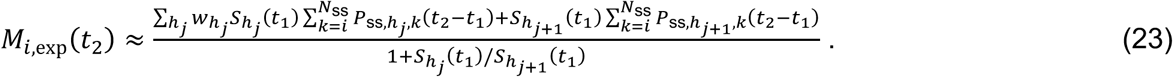

To further facilitate computation, we approximated 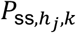 with a Γ-distribution for clone sizes *k* > 21. The Γ-distribution was parametrized by mean and variance, given by *m*(*t*) = *h*_*j*_ and σ^2^(*t*) = 2*h*_*j*_λ_ss_*t*, respectively.^63^

##### Drift of somatic variants acquired during homeostasis

We next computed the site frequency spectrum of variants acquired during homeostasis, *M*_*i*,ss_(*t*_2_) (where ‘ss’ labels variants acquired during steady state), by evaluating

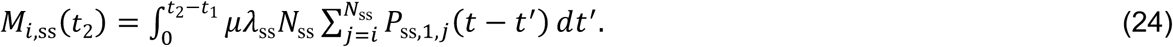

We approximated *P*_ss,1,*j*_ with a Γ-distribution if *p*(*t*)(1 − *p*(*t*)) ≥ 9 Λ *jp*(*t*)(1 − *p*(*t*)) ≥ 9. The Γ-distribution was parametrized by mean and variance, given by *m*(*t*) = 1 and σ^2^(*t*) = 2λ_ss_*t*, respectively.^63^ Moreover, if *i* ≥ 100, we approximated the sum in Eq. (24) with integration, yielding

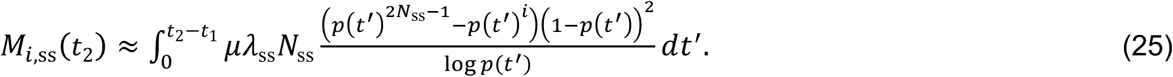

Finally, we computed the full site frequency spectrum generated by drift at *t*_2_ by summing up the variants contributed from both phases, exponential expansion (Eq. 21) and homeostasis (Eq. 25):

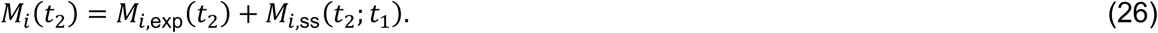

##### Frequency of somatic variants under selection

To model the site frequency spectrum of variants under selection, we first assessed the cumulative site frequency spectrum at 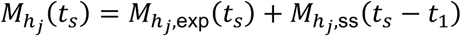 using the numerical implementation outlined above. To this end, we defined the sequence *h*_*j*_, running from clones of size 1 to *N*_ss_ as follows:

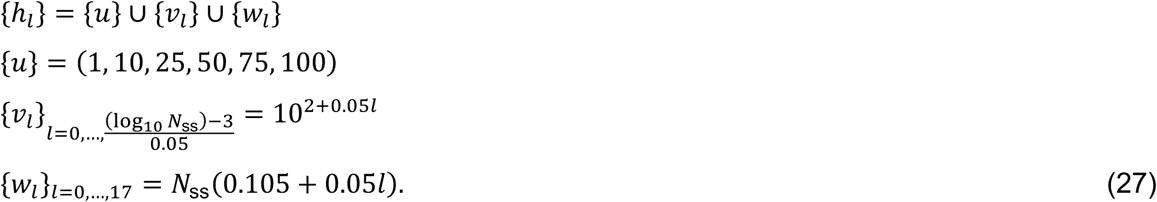

Note that we here resolved small clones at high resolution, because they may reach large frequencies due to selection. From the cumulative site frequency spectrum, we determined the histogram by computing 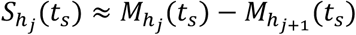. We then computed the drift of these mutations during expansion of the selected clone at both borders of each bin, and took a weighted average:

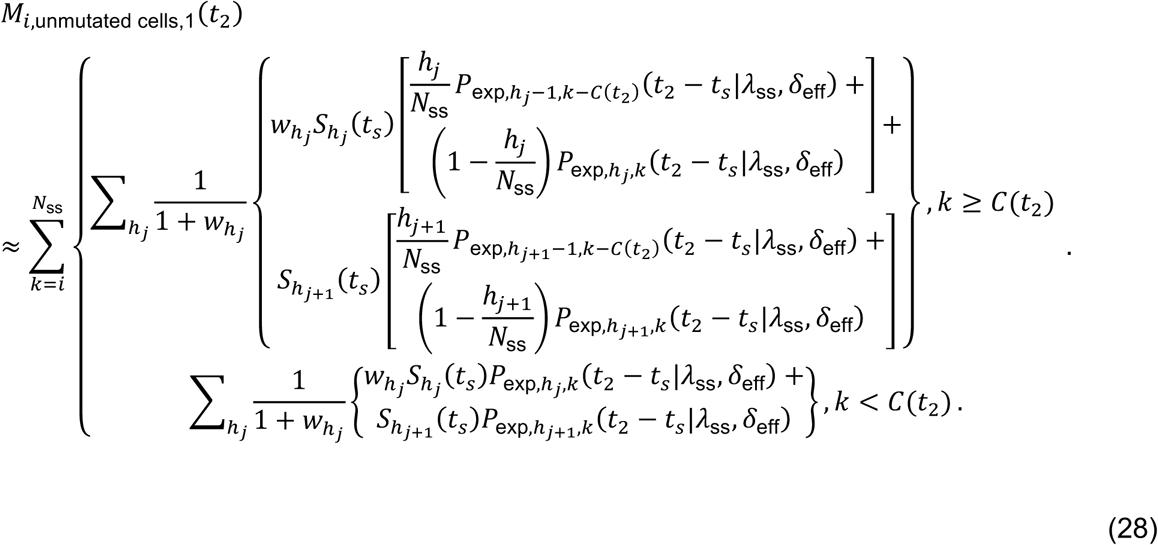

where 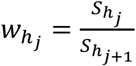 . We further facilitated computation by approximating 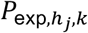 with a Γ-distribution if *h*_*j*_ + *k* > 10. The Γ-distribution was parametrized by mean and variance of *P*_exp_, given by 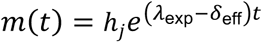 and 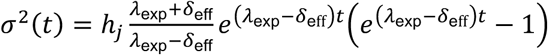, respectively.^63^

We next computed the site frequency spectrum of variants that were newly acquired during the expansion of the mutant clone by evaluating *M*_*i*,mutant cells_(*t*_2_ − t_*S*_|λ_ss_, rλ_ss_) and *M*_*i*,unmutated cells,2_ (*t*_2_ − t_*S*_|λ_ss_, *δ*_eff_) according to Eq. (9). As before, we approximated *P*_exp,a,b_ with a Γ-distribution, parametrized by mean and variance, if *a* + *b* > 10.

Finally, we computed the full site frequency spectrum at *t*_2_ by summing up the contributions of variants acquired prior to and during expansion of the CH clone:

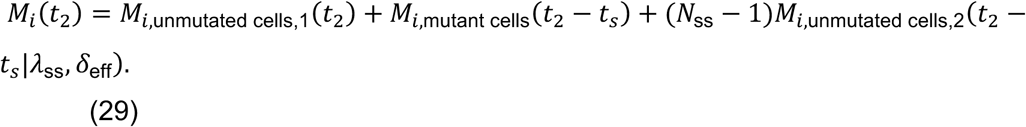

### Simulation of phylogenetic trees and in-silico evaluation of model performance

#### Stochastic simulation

To validate the population genetics model, we simulated the accumulation of variants, and the drift and selection of stem cell clones using stochastic birth-death processes. We made the functions used to simulate these processes available as part of the R package SCIFER (https://github.com/VerenaK90/SCIFER/tree/paper). During the initial expansion phase, we neglected cell loss (*δ*_exp_ = 0) and, in each simulated time step, randomly selected a cell to divide. Moreover, in order to mimic neutral variant accumulation, we introduced μ = 1 new variant in each daughter cell and memorized the mother-daughter relationship. We abrogated the expansion phase and initiated the homeostatic phase once the population reached a size of *N*_*ss*_ = 25,111 cells. Assuming that the expansion phase ends early in life, we began to track time when the system transitioned to homeostasis and defined *t* = 0 at this time point.

During the homeostatic phase, we simulated cell division and loss stochastically, using τ-leaping at constant τ = 251 (i.e., summarizing 1% of the reactions occurring in 25,000 stem cells). In each time step, we simulated τ reactions, randomly distributed to births and deaths, associated with equal probability of 0.5 due to tissue homeostasis. We first simulated the birth events by iteratively sampling random cells to divide, while introducing 1 new neutral variant per daughter cell and memorizing the phylogenetic relationships, as before. Thereafter, we simulated the death events by iteratively sampling random cells to be lost from the system. We sampled the duration of each simulated time step, d*t*, from an exponential distribution with rate *N*(*t*)2λ_*ss*_′τ, where we set λ_*ss*_ = 10/*y*ea*r*. We reported the system state at 0, 25, 50 and 75 years and terminated the simulation at 75 years.

To simulate clonal selection, we modified the simulation procedure by introducing a driver mutation with selective advantage *r* = 0.98 to a randomly selected cell at 20 years of simulation time. We stably inherited the selective advantage to the progeny of this founder cell and favored the mutant cells over normal cells for division, while assigning equal probability for them to be selected for death. To be specific, the mutant cells were selected for division with probability 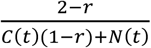, whereas normal cells were selected with probability 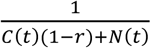, where *C*(*t*) denotes the number of mutant cells at time *t*. In case the mutant cells got randomly extinct, we re-introduced the driver in the next time step. When simulating selection we used τ = 2511 and reported the system state at 0, 25, 50 and 75 years and, in addition, once the selected clone reached clone sizes of multiples of 5%. As before, we terminated the simulation after 75 years.

#### Parameter inference

To infer the dynamic parameters, (μ, *δ*_exp_, λ_*ss*_, *N*_*ss*_, *t*_*S*_, *r*), we subsampled 10,000 cells from the simulated trees at time points specified in Supplementary Table 1 and computed the simulated VAF distribution from subsampled trees. To account for technical noise, we simulated sequencing by sampling for each variant a sequencing coverage *c* ∝ *Pois*(ĉ), where ĉ is the average sequencing coverage and thereafter sampling mutant reads according to *B*(*VAF, c*), where *B* is the binomial distribution. We simulated VAFs for coverages of 30x, 90x and 270x, and fitted the population genetics model to the simulated bulk WGS data as described above for the real data.

#### Evaluation of sensitivity and specificity

To evaluate sensitivity and specificity of out model, we analyzed the posterior probability of clonal selection. In accordance with the minimal clone sizes used in the inference setup (see above), we computed the probability of clonal selection as 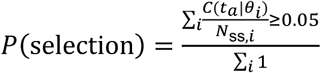 for 30x and 90x sequencing depths, and as 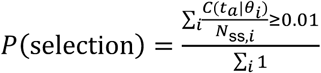for 270x sequencing depth, where *C*(*t*_*a*_|*θ*_*i*_) is the size of the selected clone at the patient age, *a*, given the *i*-th parameter sample, *θ*., and *N*_ss,*i*_ is the i-th estimate of the stem cell number. Among all parameter sets with 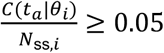 (30x and 90x coverage) or 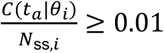 (270x coverage), we determined median clone size and rounded it with 5% accuracy if it was at least 5% or with 1% accuracy otherwise. Thereafter, we computed sensitivity and specificity of our approach, by varying *β* between 1% and 100% and classifying cases as selected if *P*(select*io*n) ≥ *β*. For each of the true clone sizes (1%, 2%, 5%, 10%, 15%, 20%, 25%, 50%, 75%; see Supplementary Table 1), we computed the number of true positives as the number of cases where the actual clone size was correctly inferred (where we classified inferred clone sizes between 0.5 and 1.5 of the actual clone size as correct). Conversely, we computed the number of false positives as the number of cases where the respective clone size was falsely inferred whereas the actual clone size was zero.

The difference between true positives and false positives was maximal for *β* = 15%. At this threshold, clones of size ≥5% VAF were reliably inferred.

### Statistics and reproducibility

Bayesian parameter inference and statistical analyses were conducted with pyABC^65^ v0.12.6, using python v3.10.1 and R (v4.2.0 and v4.2.1). We used the following R packages: ape^66^ v5.6-2, phytools^67^ v1.2-0, phangorn^68^ v2.10.0, castor^69^ v1.7.5, TreeTools v1.8.0, deSolve^70^ v1.33, openxlsx v4.2.5, cdata v1.2.0, ggpubr v0.4.0, RRphylo^71^ v2.7.0, ggplot2^72^ v3.4.2, cgwtools v3.3, ggVennDiagram v1.2.2, ggbeeswarm v0.6.0, ggsci v2.9, Hmisc v4.7.1, lemon v0.4.5, data.table v1.14.2, RColorBrewer v1.1.3, ggridges v0.5.4, doParallel v1.0.17, foreach v1.5.2, parallel v2.1, wesanderson v0.3.6, bedr v1.0.7, ggformula v0.10.2, HDInterval v0.2.2, reshape2 v1.4.4^73^, dplyr v1.0.9, scales v1.2.1.

## Data availability

bam files have been deposited on EGA (EGAS00001007558). Variant calls and model fits have been made available on Mendeley data (doi: 10.17632/yvxdb7t3yk.1). Patient information and driver mutations are available as Supplementary Data to this manuscript. If excess human DNA used in this study is available it will be made available.

## Code availability

Code to reproduce the analysis is available on https://github.com/VerenaK90/Clonal_hematopoiesis/tree/main and https://github.com/VerenaK90/SCIFER/tree/paper.

## Acknowledgments

We thank the subjects and allied health care professionals who supported collection of bone marrow and blood samples. We thank Nils Becker and all members of the Höfer and Vyas groups for discussions.

## Author contributions

V.K., N.A.J., P.V. and T.H. conceived the project. S.N., B.J.L.K., A.H.T., R.A.L., R.G., B.W., K.W., D.B., S.D. and A.J.C. collected bone marrow and blood samples. N.A.J. performed flow cytometry; N.A.J., M.M., R.M., B.U. and M.S. performed DNA extraction; N.C. prepared DNA sequencing libraries. V.K. and N.A.-P. analysed whole genome sequencing data. F.H. helped with phylogenetic inference. V.K. developed the mathematical model, performed parameter estimation and all computations. V.K., T.H. and P.V. wrote the manuscript with input from all coauthors. T.H. and P.V. acquired funding and administered the project.

## Competing interests

The authors declare no competing interests.

**Supplementary Table 7.** Prior probabilities to model site frequency spectra.

| Parameter | Description | Distribution | Lower limit | Upper limit |  |
| --- | --- | --- | --- | --- | --- |
|  |  |  |  | Bulk WGS data / simulated bulk WGS data at 30x or 90x | Pseudo-bulk WGS / simulated bulk WGS data at 270x |
| $\delta_{\text{exp}}$ | Relative loss during development | uniform | 0 | 0.75 | |
| $N_{\text{ss}}$ | Active stem cell count during homeostasis | Log-uniform | $10^{2.5}$ | $10^8$ | |
| $\mu$ | Number of neutral variants acquired per division | uniform | 0.1 | 10 | |
| $\lambda_{ss}$ | Division rate during homeostasis | Log-uniform | $10^{-3}$ | $10^{-1}$ | |
| $t_s$ | Time point of acquiring the driver | uniform | 0 | $t_a - \frac{\log 0.01 N_{ss}}{\lambda_{ss}}$ | $t_a - \frac{\log 0.001 N_{ss}}{\lambda_{ss}}$ |
| $r$ | Selective advantage associated with driver | uniform | $1 - \frac{\log N_{ss}}{\lambda_{ss}(t_a - t_s)}$ | $1 - \frac{\log 0.01 N_{ss}}{\lambda_{ss}(t_a - t_s)}$ | $1 - \frac{\log 0.001 N_{ss}}{\lambda_{ss}(t_a - t_s)}$ |
| $\Delta_{clonal}$ | Number of neutral variants present in the founder cell of the hematopoietic system | uniform | 0 | 10 | |

