## Supplementary Information for "Detecting and quantifying clonal selection in somatic stem cells"

**Supplementary Note 1. Parameter dependence of the site frequency spectrum.**

We discuss parameter inference from the measured VAF histograms, based on our theory for the site frequency spectra generated in development followed by homeostasis (Methods). Equation (8), yielding the basic site frequency spectrum (SFS) due to genetic drift, does not afford a solution in terms of elementary functions. To gain insight into how to infer the homeostatic stem cell number, $N_{\mathrm{ss}}$, the rate of self-renewing divisions, $\lambda_{\mathrm{ss}}$, and mutation rate, $\mu$, we discuss analytically solvable limiting cases and complement this by numerical calculations of the SFS. The key insights are as follows: (1) Observing neutral evolution in the homeostatic phase provides information on $N_{\mathrm{ss}}/\lambda_{\mathrm{ss}}$ (from the shape of the VAF histogram) and on $\mu N_{\mathrm{ss}}$ (from the absolute count of variants with frequency greater than a given detection threshold), both in the homeostatic phase. (2) The mutation rate $\mu$ alone shapes the SFS generated during the developmental expansion phase. The combined presence of these effects in the measured WGS data allows identifying $N_{\mathrm{ss}}$, $\lambda_{\mathrm{ss}}$ and $\mu$ separately. In particular, the time point of measurement within the transient evolution of the SFS to its invariant shape provides the necessary timescale for identifying the rate $\lambda_{\mathrm{ss}}$. (3) The appearance of a selected clone effectively contributes a second timescale, because the selected clone's age is encoded in the number of variants in its founder cell. As the clone's final size is also known, the average selected growth rate can be inferred; this information also improves inference of the stem cell parameters.

### I. Homeostatic drift

For the variants generated in homeostasis we find, inserting Equation (4) with $a = 1$ into the second integral in Equation (8), that the number of these variants with clone size *i* obeys

$$S_i(t) = \frac{\mu N_{\mathrm{ss}}}{i}\left(\frac{\lambda_{\mathrm{ss}} t}{1 + \lambda_{\mathrm{ss}} t}\right)^i . \tag{A.1}$$

Variants generated in development may start in the homeostatic phase with clone size $a > 1$; for these variants the clone size evolution cannot be given in terms of elementary functions. Nevertheless, Equation (A.1), for clones starting with size 1 in homeostasis, provides generic insight into how the site frequency spectrum in the homeostatic phase evolves as a function of the mutation rate, $\mu$, the rate of self-renewing stem cell divisions, $\lambda_{\mathrm{ss}}$, and the homeostatic number of stem cells, $N_{\mathrm{ss}}$: The number of generations, $\lambda_{\mathrm{ss}} t$, determines the shape of the SFS, while the product of mutation rate and stem cell number, $\mu N_{\mathrm{ss}}$, determines the absolute number of variants.

Equation (A.1) cannot be used directly to interpret WGS data, as the absolute clone sizes $i$ are not known in the data. Transforming to clone frequency $f = i/N_{\mathrm{ss}}$, which is measured ($f = 2\,\mathrm{VAF}$), we obtain

$$S_f(t) = \frac{\mu}{f}\left(\frac{\lambda_{\mathrm{ss}}t}{1+\lambda_{\mathrm{ss}}t}\right)^{fN_{\mathrm{ss}}}.$$

(A.2)

Expanding $\ln S_f$ about small $(\lambda_{\mathrm{ss}}t)^{-1}$, we find that the site frequency spectrum approaches

$$S_f(t) = \frac{\mu}{f}e^{-f\frac{N_{\mathrm{ss}}}{\lambda_{\mathrm{ss}}t}}$$

(A.3)

as time increases, eventually converging to $\mu/f$. The limit (A.3) is relevant for human hematopoiesis, whereas convergence to $\mu/f$ is not expected within the human lifespan (Supplementary Fig. 1a,b). As a consequence, the shape of the measured VAF histogram depends approximately on the ratio $N_{\mathrm{ss}}/\lambda_{\mathrm{ss}}$ at given time $t$. This result is reminiscent of the Fokker-Planck approximation for the clonal dynamics that contains $N_{\mathrm{ss}}/\lambda_{\mathrm{ss}}$ from the outset.[1] To additionally use the information on the absolute variant count present in the WGS data, we define the cumulative count of variants with frequency equal or greater than $f$, $M_f$,

$$M_f = \sum_{i=fN_{\mathrm{ss}}}^{N_{\mathrm{ss}}} S_i(t).$$

which is to equal $\sum\#\mathrm{variants}|_{\mathrm{VAF}\geq f/2}$ in the data. In the long-term limit, we find

$$M_f = -\mu N_{\mathrm{ss}}\ln f,$$

(A.4)

and confirm general proportionality of variant count in homeostasis with $\mu N_{\mathrm{ss}}$ by numerical calculations (Supplementary Fig. 1c).

Taken together, during neutral evolution in homeostasis the shape of the SFS as a function of VAF depends in practice on $N_{\mathrm{ss}}/\lambda_{\mathrm{ss}}$ while the absolute variant count is proportional to $\mu N_{\mathrm{ss}}$.

We show below that variants acquired during tissue development contribute information that can be used to determine the value of $\mu$.

**II. Developmental expansion**

The site frequency spectrum in a homeostatic tissue consists of variants generated during tissue homeostasis and variants generated during developmental tissue expansion. At the end of developmental expansion, $t_1$, the number of variants with frequency of at least $f$ is given by

$$\sum_{i=fN_{\mathrm{ss}}}^{N_{\mathrm{ss}}} S_i = \mu\,\lambda_{\mathrm{exp}} \sum_{i=fN_{\mathrm{ss}}}^{N_{\mathrm{ss}}} \int_0^{t_1} N(t')\,P_{\mathrm{exp},1,i}(t_1 - t')\,\mathrm{d}t'\ , \tag{A.5}$$

where $P_{\mathrm{exp},1,i}(t-t')$ is the probability to drift to a clone of size $i$ within a time span $t-t'$ according to a supercritical birth-death process (Eqs. 1 and 2). For sufficiently large expanding clones, for which drift is negligible, the cumulative site frequency spectrum approaches[2]

$$\sum_{i=fN_{\mathrm{ss}}}^{N_{\mathrm{ss}}} S_i = \frac{\mu}{1-\delta_{\mathrm{exp}}/\lambda_{\mathrm{exp}}}\frac{1}{f}. \tag{A.6}$$

Here, $\frac{\mu}{1-\delta_{\mathrm{exp}}/\lambda_{\mathrm{exp}}}$ denotes the number of mutations per an effective symmetric self-renewing stem cell division, which depends on the ratio $\delta_{\mathrm{exp}}/\lambda_{\mathrm{exp}}$ of the rates of cell loss (by differentiation and death) and proliferation. Interestingly, the posteriors of our model fits to the data yield $0 < \frac{\delta_{\mathrm{exp}}}{\lambda_{\mathrm{exp}}} < 0.5$ (Extended Data Fig. 2e,f), implying that at least half of the HSC divisions in development expand the HSC population. In turn, this implies that we can estimate the HSC mutation rate from our data with an uncertainty of about a factor of 2, as we do not know precisely the extent of HSC loss in development. Thus, at the end of expansion, the cumulative VAF distribution depends on the isolated parameter $\mu$. This information is preserved during the ensuing homeostatic phase in the VAF histogram in the range of large clone sizes.

**III. Clonal selection**

The presence of a selected subclone improves parameter inference (Eq. 18). Here, the number of variants accumulated in the subclonal cell of origin reports on the time point at which the subclone was born. The size of the selected subclone depends on its selective advantage and the stem cell number. Finally, accumulation and drift of neutral variants within the selected subclone jointly depend on the mutation rate and the growth dynamics of the selected clone (Eq. 17). In sum, subclonal selection provides a second time scale that supports inference of stem cell number and tissue dynamics.

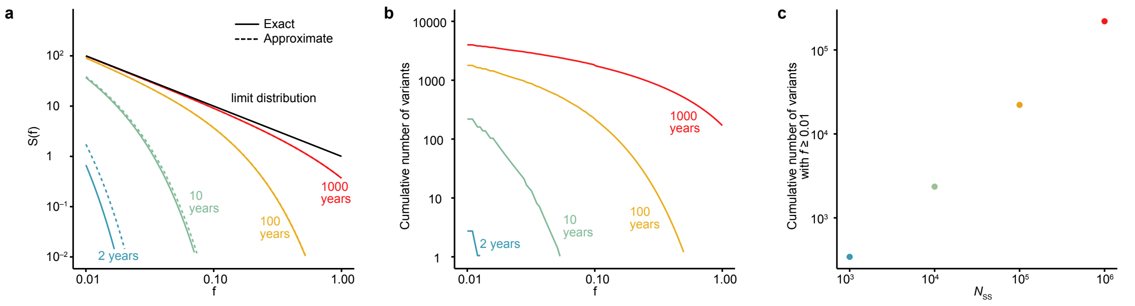


**Supplementary Fig. 1. Variant allele frequencies under genetic drift in homeostatic tissues. a,** Computed site frequency spectrum of variants acquired in a homeostatic tissue. Shown are exact (solid line, Eq. A1) and approximate solutions (dashed line, Eq. A2) at different time points along with the limit distribution for long times (black). Simulations were done with $N_{ss} = 1000$, $\lambda_{ss} = 1/\mathrm{year}$ and $\mu = 1/\mathrm{division}$. **b**, Cumulative number of variants acquired in a homeostatic tissue at different time points. Parameters are as in (**a**). **c**, Cumulative number of variants with frequency of at least 1% after 100 years for different values of $N_{ss}$ and $\lambda_{ss}$ that together yield$N_{ss}/\ \lambda_{ss} = 10^4\mathrm{years}$.


1. Watson, C. J. *et al.* The evolutionary dynamics and fitness landscape of clonal hematopoiesis. *Science* **367**, 1449-1454 (2020).
2. Bozic, I., Gerold, J. M. & Nowak, M. A. Quantifying clonal and subclonal passenger mutations in cancer evolution. *PLoS computational biology* **12**, e1004731 (2016).

**Supplementary Note 2. Peripheral blood granulocytes report on selection in HSCs.** Mononuclear cells (MNCs) are more abundant and easily accessible from peripheral blood (PB) than HSPC. However, clonal selection of T and B lymphocytes by antigens may modify the VAF distribution in this mixed population. To characterize the suitability of mature cell populations for inferring the clonal dynamics of drift and selection of HSPCs, we tested DNA from BM MNCs, BM T cell-depleted MNCs (MNC–T) and granulocytes sorted from peripheral blood for five individuals (Supplementary Table 2). We performed whole genome sequencing at 90x, applied FLORENCE to variant calls from these cell sources, and compared the results with that from $CD34^{+}$ HSPCs in the same individuals. Selection was distinguished from neutral evolution equally well in PB granulocytes and BM HSPCs (Supplementary Fig. 2). By contrast, BM MNCs gave different inference results in one individual without selection (N2), erroneously inferring a selected clone in both MNC and MNC–T samples (Supplementary Fig. 2a). In one case with selection (U3) BM MNCs erroneously missed the selected clone (Supplementary Fig. 2d). Collectively, these data suggest that selection can best be distinguished from neutral evolution in BM HSPCs and PB granulocytes, whereas MNCs should not be used.

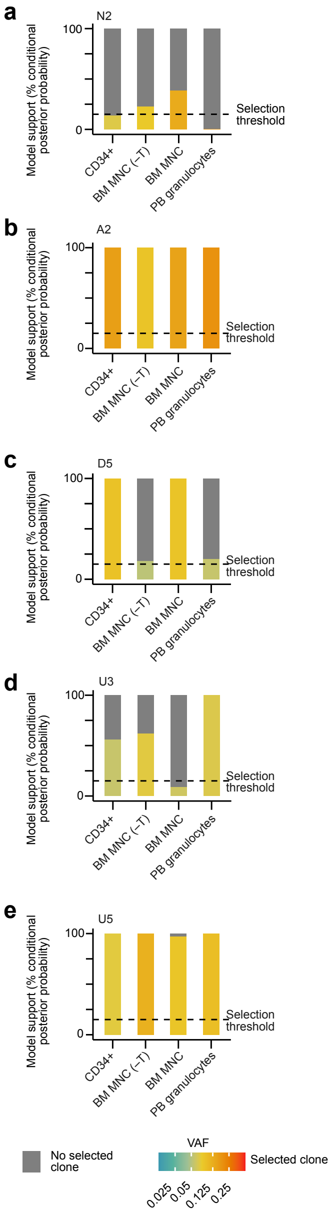


**Supplementary Fig. 2. Quantifying selection and drift in mature cell populations. a-e**, Five individuals: N2 (a), A2 (b), D5 (c), U3 (d) and U5 (e) were studied. Shown is the model support for clonal selection (posterior probability conditioned on selected clones with VAF ≥5%) and neutral evolution using DNA from CD34+ HSPC (CD34+), BM T cell depleted MNCs (BM MNC (–T)), BM MNCs and peripheral blood granulocytes. The inferred VAFs of the selected clones are color encoded; the dotted lines show the 15% selection threshold.
